# Epistasis detection and modeling for genomic selection in cowpea (*Vigna unguiculata*. L. Walp.)

**DOI:** 10.1101/576819

**Authors:** Marcus O. Olatoye, Zhenbin Hu, Peter O. Aikpokpodion

**Author notes:** Corresponding Author: Peter O. Aikpokpodion Address: Department of Genetics and Biotechnology, University of Calabar, Nigeria.

## Abstract

Genetic architecture reflects the pattern of effects and interaction of genes underling phenotypic variation. Most mapping and breeding approaches generally consider the additive part of variation but offer limited knowledge on the benefits of epistasis which explains in part the variation observed in traits. In this study, the cowpea multiparent advanced generation inter-cross (MAGIC) population was used to characterize the epistatic genetic architecture of flowering time, maturity, and seed size. In addition, considerations for epistatic genetic architecture in genomic-enabled breeding (GEB) was investigated using parametric, semi-parametric, and non-parametric genomic selection (GS) models. Our results showed that large and moderate effect sized two-way epistatic interactions underlie the traits examined. Flowering time QTL colocalized with cowpea putative orthologs of *Arabidopsis thaliana* and *Glycine max* genes like *PHYTOCLOCK1* (*PCL1* [Vigun11g157600]) and *PHYTOCHROME A* (*PHY A* [Vigun01g205500]). Flowering time adaptation to long and short photoperiod was found to be controlled by distinct and common main and epistatic loci. Parametric and semi-parametric GS models outperformed non-parametric GS model. Using known QTL as fixed effects in GS models improved prediction accuracy when traits were controlled by both large and moderate effect QTL. In general, our study demonstrated that prior understanding the genetic architecture of a trait can help make informed decisions in GEB. This is the first report to characterize epistasis and provide insights into the underpinnings of GS versus marker assisted selection in cowpea.

## Introduction

Asymmetric transgressive variation in quantitative traits is usuallly controlled by non-additive gene action known as epistasis (Rieseberg, Archer and Wayne, 1999). Epistasis has been defined as the interaction of alleles at multiple loci (Mathew *et al*., 2018). The joint effect of the alleles at these loci may be lower or higher than the total effects of these loci (Johnson, 2008). In selfing species, epistasis is common due to high level of homozygousity (Volis *et al*., 2010) and epistatic interactions have been found among loci underlying flowering time in barley (Mathew *et al*., 2018), rice (Chen *et al*., 2015; M. Chen *et al*., 2018), and sorghum (Li *et al*., 2018). However, the direct quantification of the importance of epistasis for breeding purposes has not been fully realized due to the fact that most of the current statistical models cannot efficiently characterize or account for epistasis (Mackay, 2001; Moore and Williams, 2009; Sun, Ma and Mumm, 2012; Mathew *et al*., 2018). Common quantitative traits mapping approaches are often single-locus analysis techniques. These techniques focus on the additive contribution of genomic loci (H.Barton and D.Keightley, 2002) which may explain a fraction of the genetic variation; thus leading to missing heritability.

Regardless of the limitations of genomic mapping approaches, characterization of the genetic basis of complex agronomic traits has been beneficial for breeding purposes. For example, markers tagging quantitative trait loci (QTL) have been used in marker-assisted selection (MAS) in breeding programs (Zhang *et al*. 2003; Pan *et al*. 2006; Saghai Maroof *et al*. 2008; Foolad and Panthee 2012; Massman *et al*. 2013; Mohamed *et al*. 2014; Zhao *et al*. 2014). However, the efficiency of QTL based MAS approach in breeding is limited. First, the small sample size of bi-parental populations where QTL are detected often result in overestimation of the respective QTL effect sizes; a phenomenon known as Beavis effect (Utz, Melchinger and Schön, 2000; Xu, 2003; King and Long, 2017). Second, genetic diversity is limited to the two parents forming the bi-parental population, thus QTL may not reflect the entire variation responsible for the trait and may not be transferable to other genetic backgrounds (Xu *et al*., 2017). Third, linkage mapping is limited in power to detect small effect loci, thus only the available large effects loci are used for MAS (Ben-Ari and Lavi 2012). Notably, MAS is more efficient with traits controlled by few genomic loci and not polygenic traits (Bernardo, 2008). In contrast, genomic selection (GS) that employs genome wide markers has been found to be more suited for complex traits, and also having higher response to selection than MAS (Bernardo and Yu, 2007; Wong and Bernardo, 2008; Cerrudo *et al*., 2018).

In GS, a set of genotyped and phenotyped individuals are first used to train a model that estimates the genomic estimated breeding values (GEBVs) of un-phenotyped but genotyped individuals (Jannink, Lorenz and Iwata, 2010). GS models often vary in performance with the genetic architecture of traits. Parametric GS models are known to capture additive genetic effects but not efficient with epistatic effects due to the computational burden of high-order interactions (Moore and Williams, 2009; Howard, Carriquiry and Beavis, 2014). Parametric GS models with incorporated kernels (marker based relationship matrix) for epistasis have recently been developed (Covarrubias-pazaran, 2017). Semi-parametric and non-parametric GS models capturing epistatic interactions have been developed and implemented in plant breeding (Gianola, Fernando and Stella, 2006; Gianola and de los Campos, 2008; De Los Campos *et al*., 2010). Semi-parametric models as Reproducing Kernel Hilbert Space (RKHS) reduces parametric space dimensions to efficiently capture epistatic interactions among markers (Jiang and Reif, 2015; de Oliveira Couto *et al*., 2017). Using simulated data, Howard *et al*. 2014 showed that semi-parametric and non-parametric GS models can improve prediction accuracies under epistatic genetic architectures. In general, GS has been widely studied in and applied to major crop species including both cereals and legumes. However, in orphan crop species, applications of genomic-enabled breeding (GEB) methods is still limited (Varshney *et al*., 2012).

Cowpea (*Vigna unguiculata* L. Walp) is a widely adapted warm-season orphan herbaceous leguminous annual crop and an important source of protein in developing countries (Muchero *et al*., 2009; Varshney *et al*., 2012; Boukar *et al*., 2018; Huynh *et al*., 2018). Cowpea is cultivated over 12.5 million hectares in tropical and sub-tropical zones of the world including Sub-Saharan Africa, Asia, South America, Central America, the Caribbean, United States of America and around the Mediterranean Sea. However, more than 95 *per cent* of cultivation takes place in Sub-Saharan Africa (Boukar *et al*., 2018). It is the most economically important African leguminous crop and of vital importance to the livelihood of several millions of people. Due to its flexibility as a “hungry season crop” (Langyintuo *et al*., 2003), cowpea is part of the rural families’ coping strategies to mitigate the effect of changing climatic conditions.

Cowpea’s nitrogen fixing and drought tolerance capabilities make it a valuable crop for low-input and smallholder farming systems (Hall *et al*., 2003; Boukar *et al*., 2018). Breeding efforts using classical approaches have been made to improve cowpea’s tolerance to both biotic (disease and pest) and abiotic (drought and heat) stressors (Hall *et al*., 2003; Hall, 2004). Advances in applications of next generation sequencing (NGS) and development of genomic resources (consensus map, draft genome, and multiparent population) in cowpea have provided the opportunity for the exploration for GEB (Muchero *et al*., 2009; Boukar *et al*., 2018; Huynh *et al*., 2018). MAS and GS have improved genetic gain in soybean (*Glycine max*) (Jarquin, Specht and Lorenz, 2016; Kurek, 2018; Matei *et al*., 2018) and common bean (*Phaseolus vulgaris*) (Schneider, Brothers and Kelly, 1997; Yu, Park and Poysa, 2000; Wen *et al*., 2019). However, cowpea still lags behind major legumes in the area of GEB applications. GEB has the potential of enabling expedited cowpea breeding to ensure food security in developing countries where national breeding programs still depend on labor-intensive and time-consuming classical breeding approaches.

In this study, the cowpea multiparent advanced generation inter-cross (MAGIC) population was used to explore MAS and GS. The cowpea MAGIC population was derived from intercrossing among eight founder lines (Huynh *et al*., 2018) and offers greater genetic diversity than bi-parentals to identify higher-order epistatic interactions (Mathew *et al*., 2018). Although, theoretical models and empirical studies involving simulations have suggested the significant role for epistasis in breeding (Melchinger *et al*., 2007; Volis *et al*., 2010; Messina *et al*., 2011; Howard, Carriquiry and Beavis, 2014); empirical evidence from practical breeding are limited. Therefore, the epistatic genetic architecture of three traits in cowpea was evaluated alongside its considerations in genomic enabled breeding using parametric, semi-parametric, and non-parametric GS models.

## Materials and Methods

### Plant genetic resource and phenotypic evaluation

This study was performed using publicly available cowpea MAGIC population’s phenotypic and genotypic data (Huynh *et al*., 2018). The MAGIC population was derived from intercross between eight founders. The F_1s_ were derived from eight-way intercross between the founders and were subsequently selfed through single seed descent for eight generations. The F_8_ RILs were later genotyped with 51,128 SNPs using the Illumina Cowpea Consortium Array. A core set of 305 MAGIC RILs were selected and phenotyped (Huynh *et al*., 2018). The RILs were evaluated under two irrigation regimes.

In this study, the flowering time (FLT), maturity (MAT), and seed size (SS) data were analyzed for environment-by-environment correlations and best linear unbiased predictions (BLUPs). The traits analyzed in this study are; FTFILD (flowering time under full irrigation and long day), FTRILD (flowering time under restricted irrigation and long day), FTFISD (flowering time under full irrigation and short day), FTRISD (flowering time under restricted irrigation and short day), FLT_BLUP (BLUP of flowering time across environments), MFISD (maturity under full irrigation and short day), MRISD (maturity under restricted irrigation and short day), MAT_BLUP (BLUP of maturity across environments), SSFISD (seed size under full irrigation and short day), SSRISD (seed size under restricted irrigation and short day), SS_BLUP(BLUP of seed size across environments). In addition, using both genomic and phenotypic data, narrow sense heritability was estimated using RRBLUP package in R (Endelman, 2011).

### QTL and Epistasis Mapping

QTL mapping was performed for all traits using the stepwise regression model implemented in TASSEL 5.0 standalone version (Bradbury *et al*., 2007). The approach implements both forward inclusion and backward elimination steps. The model accounts for major effect loci and reduces collinearity among markers. The model was designed for multi-parental populations and no family term was used in the model since MAGIC population development involved several steps of intercross that reshuffles the genome and minimizes phenotype-genotype covariance. A total of 32,130 SNPs across 305 RILs were used in the analysis. A permutation of 1000 was used in the analysis.

To characterize the epistatic genetic architecture underlying flowering time, maturity, and seed size, the Stepwise Procedure for constructing an Additive and Epistatic Multi-Locus model (SPAEML; (Chen *et al*., 2018)) epistasis pipeline implemented in TASSEL 5.0 was used to perform epistasis mapping for phenotypic traits (FTFILD, FTRILD, FTFISD, FTRISD, FT_BLUP, MFISD, MRISD, MT_BLUP, SSFISD, SSRISD, and SS_BLUP). One critical advantage of SPAEML that led to its consideration for this study is its ability to correctly distinguish between additive and epistatic QTL. SPAEML source code is available at https://bitbucket.org/wdmetcalf/tassel-5-threaded-model-fitter. The minor allele frequency of each QTL was estimated using a custom R script from http://evachan.org/rscripts.html. The proportion of phenotypic variation explained (PVE) by each QTL from both QTL and Epistasis mapping was estimated by multiplying the *R*^2^ obtained from fitting a regression between the QTL and the trait of interest by 100. The regression model for estimating PVE is;

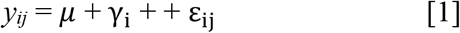

where *y_ij_* is the phenotype, *μ* is the overall mean, γ_i_ is the term for QTL, and ε_*ij*_ is the residual term.

A set of *a priori* genes (n=100; Data S1) was developed from *Arabidopsis thaliana* and *Glycine max* flowering time and seed size genes obtained from literature and https://www.mpipz.mpg.de/14637/Arabidopsis_flowering_genes. The cowpea orthologs of these genes were obtained by blasting the *A. thaliana* and *G. max* sequence of the *a priori* genes on the new *Vigna* genome assembly *v.1* on Phytozome (Goodstein *et al*., 2012). The corresponding cowpea gene with the highest score was selected as a putative ortholog. Colocalizations between the cowpea putative orthologs and QTL were identified using a custom R script.

### Marker Assisted Selection Pipeline

In order to evaluate the performance of MAS in cowpea, a custom pipeline was developed in R. First, using subbagging approach, 80% of the 305 RILs randomly sampled without replacement was used as the training population; followed by performing a Multi-locus GWAS (Multi-locus Mixed Model, MLMM) (Segura *et al*., 2012) on both genomic and phenotypic data of the training population. The MLMM approach implements stepwise regression involving both forward and backward regressions. This model accounts for major effect loci and reduces the effect of allelic heterogeneity. A K-only model that accounts for a random polygenic term (kinship relationship matrix) was used in the MLMM model. No term for population structure was used in the model since MAGIC population development involved several steps of intercross that reshuffles the genome and minimizes phenotype-genotype covariance. A total of 32130 SNPs across 305 RILs were used in the GWAS analysis and coded as −1 and 1 for homozygous SNPs and 0 for heterozygous SNPs. Bonferroni correction with α = 0.05 was used to determine the cut-off threshold for each trait association (α/total number of markers = 1.6 e-06).

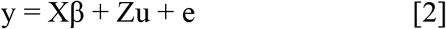

where y is the vector of phenotypic data, β is a vector of fixed effects other than SNP or population structure effects; u is an unknown vector of random additive genetic effects from multiple background QTL for RILs. X and Z are incident matrices of 1s and 0s relating y to β and u (Yu *et al*., 2006).

Second, top three most significant associations were then selected from the genomic data of the training population to train a regression model by fitting the SNPs in a regression analysis with the phenotypic information. This training model was later used alongside the predict function in R to predict the phenotypic information of the validation population (20% that remained after sub-setting the training population). The prediction accuracy of MAS was obtained as the correlation between this predicted phenotypic information and the observed phenotypic information for the validation data.

### Genomic Selection Pipeline

In order to evaluate the performance of using known QTL as fixed effects in GS models and to compare the performance of parametric, semi-parametric and non-parametric GS models; a custom GS pipeline was developed in R. The GS pipeline was made up of four GS models, which were named as FxRRBLUP (Ridge Regression BLUP where markers were fitted as both fixed and random effects; parametric), RRBLUP (RRBLUP where markers were only fitted as random effects; parametric), Reproducing Kernel Hilbert Space (RKHS; semi-parametric), and Support Vector Regression (SVR; non-parametric). First, using subagging approach, 80% of the RILs were randomly sampled without replacement (training population) followed by running MLMM GWAS and selecting the three most significant associations, which were used as fixed effects in the FxRRBLUP. These three SNPs were removed from the rest of SNPs that were fitted as random effects in the FxRRBLUP model. The RRBLUP, RKHS, and SVR models were fitted simultaneously in the same cycle as FxRRBLUP to ensure unbiased comparison of GS models. Likewise, in order to ensure unbiased comparison between GS and MAS approaches; similar seed numbers were used for the subagging sampling of training populations across 100 cycles for GS and MAS. The validation set was composed of the remaining 20% of the RILs after sampling the 80% (training set). Prediction accuracy in GS was estimated as the Pearson correlation between measured phenotype and genomic estimated breeding values of the validation population. Also, for flowering time, each environment was used as a training population to predict the other three environments.

### Ridge Regression BLUP (RRBLUP)

The RRBLUP models without and with fixed effects can be described as;

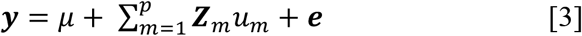

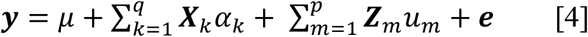

where **y** is the vector (*n* x 1) of observations (simulated phenotypic data), *μ* is the vector of the general mean, *q* is the number of selected significant associated markers (q=3), ***X_k_*** is the *k*^h^ column of the design matrix ***X***, *α* is the fixed additive effect associated with markers *k … q, **u*** random effects term, with 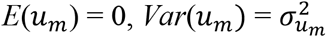 (variance of marker effect), *p* is the marker number (*p* > *n*), ***Z_m_*** is the *m*^th^ column of the design matrix ***Z***, *u* is the vector of random marker effects associated with markers *m* … *p*. In the model, ***u*** random effects term, with 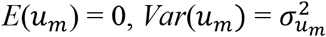 (variance of marker effect), *Var*(**e**) = *σ*^2^ (residual variance), *Cov*(u, **e**) = 0, and the ridge parameter *λ* equals 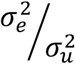 (Meuwissen, Hayes and Goddard, 2001; Endelman, 2011; Howard, Carriquiry and Beavis, 2014). In this study RRBLUP with and without fixed effects were implemented using the *mixed.solve* function in *rrBLUP* R package (Endelman, 2011).

### Reproducing Kernel Hilbert Space (RKHS)

Semi-parametric models are known to capture interactions among loci. The semi-parametric GS approach used in this study was implemented as Bayesian RKHS in *BLGR* package in R (Perez, 2014), and described as follows:

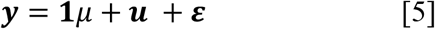

where ***y*** is the vector of phenotype; **1** is a vector of 1’s; *μ* is the mean; ***u*** is vector of random effects ~MVN 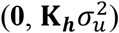; and ***ε*** is the random residual vector ~ MVN 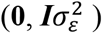. In Bayesian RKHS, the priors *p*(*μ, **u, ε***) are proportional to the product of density functions MVN 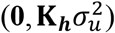 and MVN 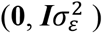. The kernel entries matrix (**K_*h*_**) with a Gaussian kernel uses the squared Euclidean distance between marker genotypes to estimate the degree of relatedness between individuals, and a smoothing parameter (*h*) multiplies each entry in **K_*h*_** by a constant. In the implementation of RKHS a default smoothing parameter *h* of 0.5 was used alongside 1,000 burns and 2,500 iterations.

### Support Vector Regression (SVR)

Support vector regression method (Vapnik, 1995; Maenhout *et al*., 2007; Long *et al*., 2011) was used to implement non-parametric GS approach in this study. The aim of the SVR method is to minimize prediction error by implementing models that minimizes large residuals (Long *et al*., 2011). Thus, it is also referred to as the “*ε*-intensive” method. It was implemented in this study using the normal radial function kernel (*rbfdot*) in the *ksvm* function of *kernlab* R package (Karatzoglou *et al*., 2004).

### Parameters evaluated in GS and MAS

Additional parameters were estimated to further evaluate the performance of GS and MAS models. A regression model was fitted between observed phenotypic information and GEBV of the validation set to obtain both intercept and slope for both GS and MAS in each cycle of prediction. The estimates of the intercept and slope of the regression of the observed phenotypic information on GEBVs are valuable since their deviations from expected values can provide insight into deficiencies in the GS and MAS models (Daetwyler *et al*., 2013). The bias estimate (slope and intercept) signify how the range of values in measured and predicted traits differ from each other. In addition, the coincidence index between the observed and GEBVs for both GS and MAS models was evaluated. The coincidence index (Fernandes *et al*., 2018) evaluates the proportion of individuals with highest trait values (20%) that overlapped between the measured phenotypes and predicted phenotypic trait values for the validation population.

### Evaluation of the effect of marker density and training population size

The effect of marker density and training population size on GS performance were evaluated. GS was performed using 20% (6426 SNPs), 40% (12852), 60% (19278), and 80% (25704) of the total number of markers available in this study (32130). Each proportion of the aforementioned marker densities was randomly sampled without replacement and used for training GS models and predict in the validation set and repeated for 100 cycles. Furthermore, to evaluate the effect of training population size on prediction accuracy, four levels (20% (61 RILs), 40% (122 RILs), 60% (183 RILs), and 80% (244 RILs)) of total population size (305 RILs) were used to train GS model and validate only 20% (61 RILs) of the total population size (305 RILs). Subagging approach was used to subset the training and validation sets at a time and repeated for 100 cycles.

## Results

### Phenotypic and genotypic variation in cowpea

Results showed variation between number of days to 50% flowering under long-day photoperiod and short-day photoperiod. Days to flowering time were higher for RILs under long-day than short-day (Figure 1). Results showed positive correlations between environments for each trait (Table S1 and S2). Furthermore, genomic heritability were moderate for the traits ranging between 0.41 under long day photoperiod to 0.48 for flowering time under short-day photoperiod, 0.21 under restricted irrigation to 0.30 under full irrigation for maturity, and 0.39 under restricted irrigation to 0.47 under full irrigation for seed size (Table S1 and S2).

**Figure 1:**
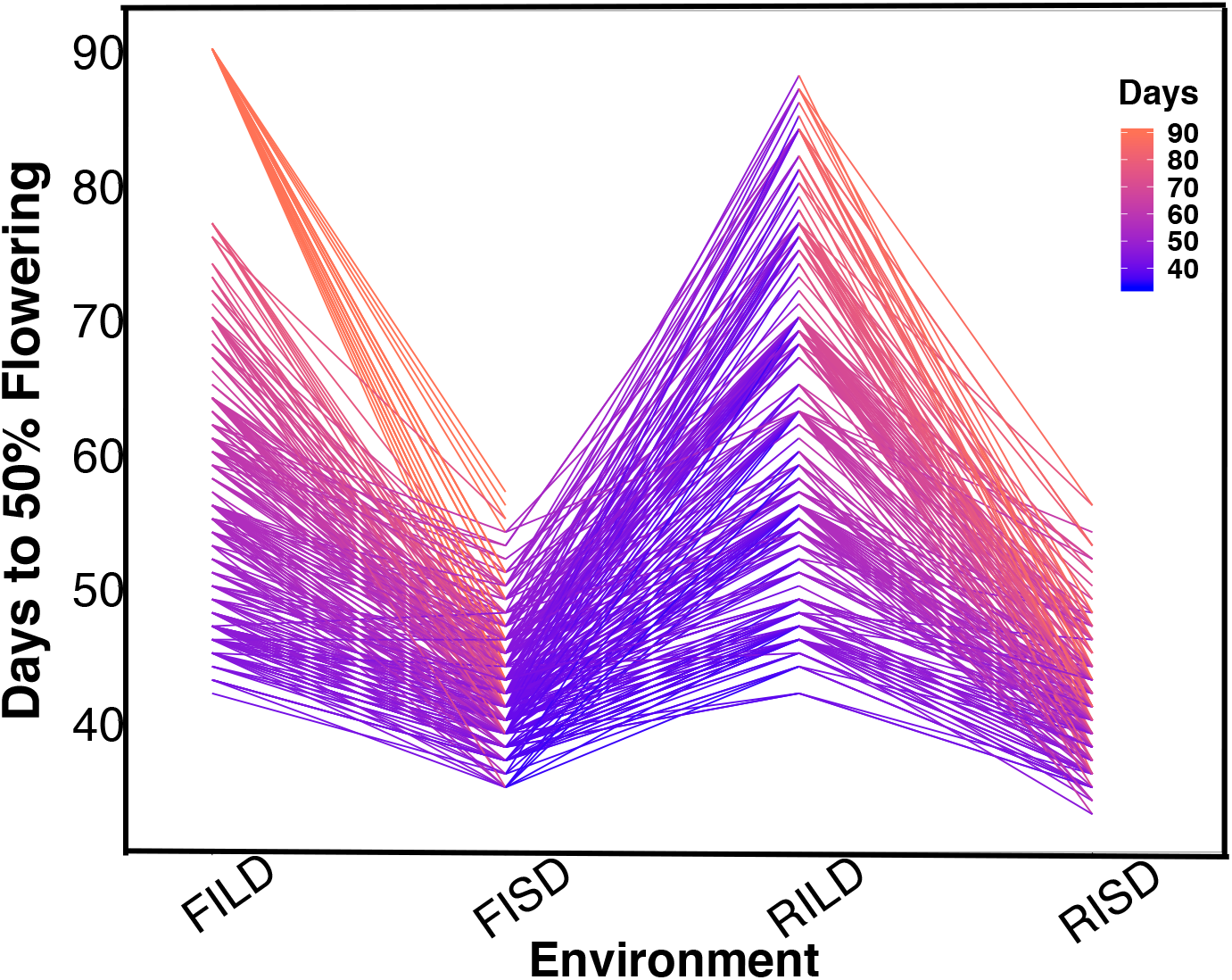
The norm of reaction plot for flowering time variation under long-day and short-day periods. Evaluation environments are represented on the x-axis (full irrigation and long day [FILD], full irrigation and short day [FISD], restricted irrigation and long day [RILD], and restricted irrigation and short day [RISD]). The number of days to 50% flowering is represented on the y-axis.

### Genetic architecture of traits

#### Main effect QTL

The cowpea MAGIC population facilitated the characterization of the genetic architecture of flowering time, maturity and seed size. In this study QTL associated with flowering time, maturity, and seed size were identified using stepwise regression analysis (Table S3, Data S2). Results showed that 32 QTL (22 unique) in total were associated with flowering time traits (FT_BLUP [8 QTL, explaining 73.2 % of phenotypic variation (PV)], FTFILD [5 QTL, explaining 66.2% of PV], FTRILD [5 QTL explaining 48.6% of PV], FTFISD [8 QTL explaining 52.1% of PV], and FTRISD [6 QTL explaining 43.9% of PV]). Each of the total QTL associated with flowering time traits explained between 2% to 28% of the phenotypic variation. QTL qVu9:23.36, qVu9:24.77, and qVu9:22.65 (MAF= 0.29, 0.28, and 0.49) explained the largest proportion of variation (28%, 24%, and 19%) with additive effects of 7, 7, and 6 days respectively. The minor allele frequency (MAF) of the flowering time QTL ranges from 0.13 to 0.50. For maturity traits, 13 QTL (11 unique QTL) in total were identified with five QTL (explaining 35.9% of PV) for MAT_BLUP, 4 QTL (explaining 24.5% of PV) for MFISD, and 4 QTL (explaining 27.9% of PV) for MRISD. All maturity traits QTL explained between 4.5 to 10% of phenotypic variation and MAF ranges from 0.15 to 0.49.

Furthermore, for seed size traits, 10 QTL (7 unique QTL) in total were identified with 3 QTL (explaining 39.3% of PV) for SS_BLUP, 3 QTL (explaining 41% of PV) for SSFISD, and 4 QTL (explaining 39.4% of PV) for SSRISD. QTL qVu8:74.21, qVu8:74.29, qVu8:76.81 associated with SSFISD, SS_BLUP, and SSRISD explained the largest PV (29%, 25%, and 20%). All seed size trait QTL explained between 3 to 29% of PV and with MAF range between 0.21 and 0.49. A pleiotropic QTL qVu8:74.21 (MAF=0.24) was associated with both MRISD and SSRISD (explained 5% and 29% of PV respectively).

#### Two-way epistatic interaction QTL

Currently there is limited knowledge about what role epistasis plays in phenotypic variation in cowpea. Our results identified epistatic QTL underlying flowering time, maturity, and seed size (Table S4, Data S3). For flowering time traits, there were 42 two-way epistatic interactions at 84 epistatic loci (only 65 loci were unique). Among these are; 20 epistatic loci for FLT_BLUP, 18 epistatic or FTFILD, 12 epistatic loci for FTRILD, 14 epistatic loci for FTFISD, and 20 epistatic loci for FTRISD. Some large effect loci were involved in epistatic interactions in flowering time; examples include, QTL qVu9:25.39 (MAF=0.28, FT_BLUP PVE=23.5%, FTFILD PVE=24.5%, FTRILD PVE=26%) and QTL qVu9:3.46 (MAF=0.35, FLT_BLUP PVE=13.5%, FTRILD PVE=14.1%). For maturity, there were 17 pairwise epistatic interactions across 34 loci (of which 30 were unique). Among the maturity QTL, qVu9:8.37 had the largest effect explaining ~9% of the phenotypic variation. One epistatic interaction overlapped with both FTRISD, MRISD, and MT_BLUP (qVu2:48.05+ qVu9:8.37, MAF=0.30 and 0.39 respectively). For seed size, there were 13 interactions at 26 loci (19 were unique). Only one QTL (qVu8:74.29, MAF=0.25) had interactions with multiple QTL. The largest effect epistatic QTL associated with the three seed size traits (SS_BLUP, SSFISD, and SSRISD) is qVu8:74.29 (MAF0.25). Some QTL were found to overlap among main effect QTL and epistatic effect QTL for flowering time (nine QTL), maturity (three QTL), and seed size (three QTL) (Figure S1).

#### Main effect and epistatic QTL colocalized with *a priori* genes

Gene functions can be conserved across species (Huang *et al*., 2017). In this study, a set of a priori genes was compiled from both *A. thaliana* and *G. max*. Both main effect QTL and epistatic QTL colocalized with putative cowpea orthologs of *A. thaliana* and *G. max* flowering time and seed size genes (Figure 2 – 5, Figure S2 – S11, Data S4). A putative cowpea ortholog (Vigun09g050600) of *A. thaliana* circadian clock gene *phytochrome E* (*PHYE*; AT4G18130) (Aukerman and Sakai, 2003) colocalized with FTFILD QTL (qVu9:22.65; PVE=19.5%; main effect QTL) at the same genetic position. Also, a putative cowpea ortholog (Vigun07g241700) of *A. thaliana* circadian clock gene *TIME FOR COFFEE* (*TIC*; AT3G22380) (Hall *et al*., 2003) colocalized at the same genetic position with FTFISD QTL (qVu7:86.92; PVE=2.6%; main effect QTL). The cowpea flowering time gene (*VuFT*; Vigun06g014600; CowpeaMine *v*.06) colocalized with an epistatic QTL (qVu6:0.68; PVE=3.5%) associated with FLT_BLUP and FTRILD at the same genetic position. The cowpea ortholog (Vigun11g157600) of *A. thaliana* circadian clock gene *PHYTOCLOCK1* (*PCL1*; AT3G46640) (Hazen *et al*., 2005) colocalized with an epistatic QTL (qVu11:50.94; PVE=8-10%) associated with both FTFILD and FTRILD at the same genetic position. A putative cowpea ortholog (Vigun11g148700) of *A. thaliana* photoperiod gene *TARGET OF EAT2* (*TOE2*; AT5G60120) (Mathieu *et al*., 2009) was found at a proximity of 0.6cM from a QTL (qVu11:49.06; PVE=7-11%; main effect QTL) associated with FTFILD, FTRILD, and FLT_BLUP. Some of the *a priori* genes colocalized with some QTL that are both main effect and epistatic QTL. For instance, the cowpea ortholog (Vigun01g205500) of *G. max* flowering time gene *phytochrome A* (*PHYA*; Glyma19g41210) (Tardivel *et al*., 2014) colocalized with a FTFILD QTL (qVu1:66.57; PVE=5.3%; both main effect and epistatic QTL) at the same genetic position (Data S4). Lastly, a putative cowpea ortholog (Vigun08g217000) of *A. thaliana histidine kinase2* gene (*AHK2*; AT5G35750) (Orozco-Arroyo *et al*., 2015) was found at a proximity of about 1-2cM from three QTL (qVu8:74.29, qVu8:74.21, qVu8:76.81; PVE=25%, 29.3%, and 20% respectively; main effect and epistatic QTL) associated with seed size traits SS_BLUP, SSFISD, and SSRISD).

**Figure 2:**
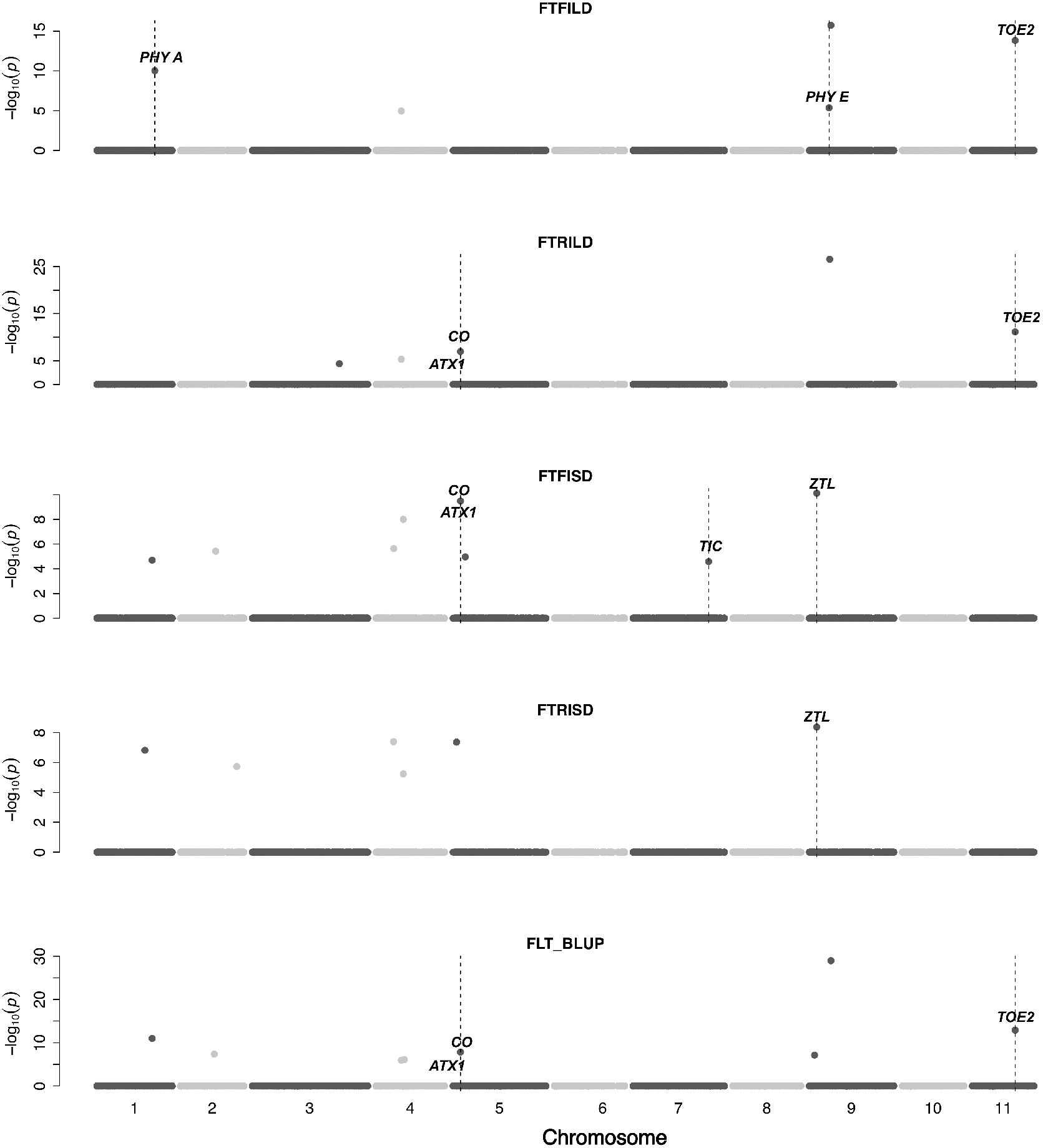
Main QTL plot for flowering time traits in the cowpea MAGIC population. QTL plots for flowering time under full irrigation and long day (FTFILD), flowering time under restricted irrigation and long day (FTRILD), flowering time under full irrigation and short day (FTFISD), flowering time under restricted irrigation and short day (FTRISD), and BLUPs of environments (FLT_BLUP). The chromosome numbers are located on the x-axis and the negative log of the *P*-values on the y-axis. The genetic position of the colocalization between QTL and *a priori* genes are indicated by broken vertical lines. The texts displayed on the vertical broken lines are the names of *a priori genes*.

In addition, some a priori genes were associated with multiple traits. The putative cowpea ortholog (Vigun05g024400) of *A. thaliana* circadian clock gene *CONSTANS* (*CO*; AT5G15840) (Wenkel *et al*., 2006) colocalized at the same genetic position with a QTL (qVu5:8.5; PVE=6-8%; both main effect and epistatic QTL) associated with flowering time and maturity traits (FLT_BLUP, FTFISD, FTRILD, FTRISD, MAT_BLUP, and MFISD); The putative cowpea ortholog (Vigun09g025800) of *A. thaliana* circadian clock gene *ZEITLUPE* (*ZTL*; AT5G57360) (Somers *et al*., 2000) colocalized at the same genetic position with a QTL (qVu9:8.37; PVE=9-11%; both main effect and epistatic QTL) associated with flowering time and maturity traits (FTFISD, FTRISD, and MRISD).

#### GS and MAS for flowering time

Prior knowledge about the genetic architecture of a trait can help make informed decisions in breeding. First to compare the performance of GS and MAS models for flowering time within each daylength results showed that under long day length (FTFILD and FTRILD); FxRRBLUP (mean prediction accuracy [mPA] = 0.68, 0.68; mean coincidence index [mCI]=0.49, 0.40) and MAS [mPA=0.64, 0.61; mCI=0.45, 0.37] outperformed RRBLUP [mPA=0.55, 0.58; mCI=0.37, 0.35], RKHS [mPA=0.55, 0.58; mCI=0.37, 0.36], and SVR [mPA=0.54, 0.50; mCI=0.35, 0.28] (Figures 6 and 7, Table S 3 and 4). For flowering time under long day, coincidence index values were higher under full irrigation than under restricted irrigation. For flowering time under short day (FTFISD and FTRISD), all GS models outperformed MAS [mPA=0.33, 0.25; mCI=0.30, 0.26]. Among the GS models, RKHS and RRBLUP had the highest prediction accuracies. However, the coincidence index of FxRRBLUP was higher than the rest of the GS models for FTRISD. In general, the mean of the slope and intercept for the GS models except SVR were usually close to the expected (1 and 0) (Figure S12–S13). MAS also deviated away from the expected slope and intercept (1 and 0) more than the FxRRBLUP, RKHS, and RRBLUP for FTRISD (Figure S12–S13). Second to evaluate the effect of photoperiod and irrigation regime on the performance of training population, each environment (day length and irrigation regime combination) was used as a training population to predict the rest in a di-allele manner. Results showed that prediction accuracy between environments in the same photoperiod was higher than environments in different photoperiod (Figure S14). Also, when training populations were under full irrigation, their prediction accuracies were higher than when training populations were under restricted irrigation (Figure S14). For FT_BLUP, GS models outperformed MAS except SVR which had the same mPA [0.59] as MAS while FxRRBLUP had the highest mPA and mCIs among the GS models (Figure S 15 and 16).

#### GS and MAS for maturity and seed size

For maturity (MT_BLUP, MFISD, and MRISD), RKHS and RRBLUP had better performance (Figures 6 and 7; Table S4 and S5) than the rest of the models including MAS. All models deviated from the expected slope and intercept estimates, but RRBLUP had the least deviation for MRISD. For seed size, FxRRBLUP had the best performance followed by MAS compared to the rest of the GS models (RKHS, RRBLUP, and SVR) (Figures 6 and 7; Table S5 and S6). GS and MAS models had varying levels of deviation from the expected estimates of slope and intercept. RKHS and RRBLUP were closer to the expected than FxRRBLUP and MAS (Figure S12–S13). SVR had the highest deviation.

#### Effect of marker density and training population size

The effect of marker density and population size on GS in cowpea was investigated with the aim of making recommendations for resource limited national research centers in developing countries. For the effect of marker density on prediction accuracy, no significant relationship was observed between marker densities for MTBLUP while a significant increase in prediction accuracy was only observed between marker density 20% - 60% for FTBLUP, and between marker densities 40% - 60% and 40% - 80% for SSBLUP (Figure S19A). For the training population size effect, results revealed that prediction accuracy increased with increasing the size of the training set. All difference between training set sizes were significantly increased with the training population size increase (Tukey test *P*-value < 0.001) (Figure S19B).

## Discussion

### Epistasis play important roles in determining the genetic architecture of agronomic traits in cowpea

Multi-parental populations have demonstrated ability to facilitate robust characterization of genetic architecture in terms of genetic effect size, pleiotropy, and epistasis (Buckler *et al*., 2009; Brown *et al*., 2011; Peiffer *et al*., 2014; Bouchet *et al*., 2017; Mathew *et al*., 2018). Using the cowpea MAGIC population, this study showed that both additive main QTL and additive x additive epistatic QTL with large and (or) moderate effects underlie flowering time, maturity, and seed size in cowpea. Although most of the epistatic QTL identified were two-way interacting loci, results showed some of them were involved in interactions with more than one independent loci (Figure 3–5 and Figure S4–11). This implies the possibility of three-way epistatic interactions underlying some of the traits. Our inability to identify and discuss three-way epistatic interactions is due to the mapping approach used, which only mapped two-way epistatic interactions. Three-way epistatic interactions have been found to underlie flowering time in the selfing crop specie barley (Mathew *et al*., 2018). Furthermore, overlaps between main and epistatic QTL (Figure S2) indicate these to be main QTL that are involved in epistatic interactions with other loci. However, one caveat that may also be responsible for some of the QTL among the overlaps is the false positive rate of SPEAML. The SPEAML software used for epistasis mapping showed high false positive rate with a sample size of 300 individuals (Chen *et al*., 2018). It is possible that some of the overlapped QTL are main QTL that were miscategorized as epistatic loci by SPEAML since our cowpea MAGIC population had 305 RILs.

**Figure 3:**
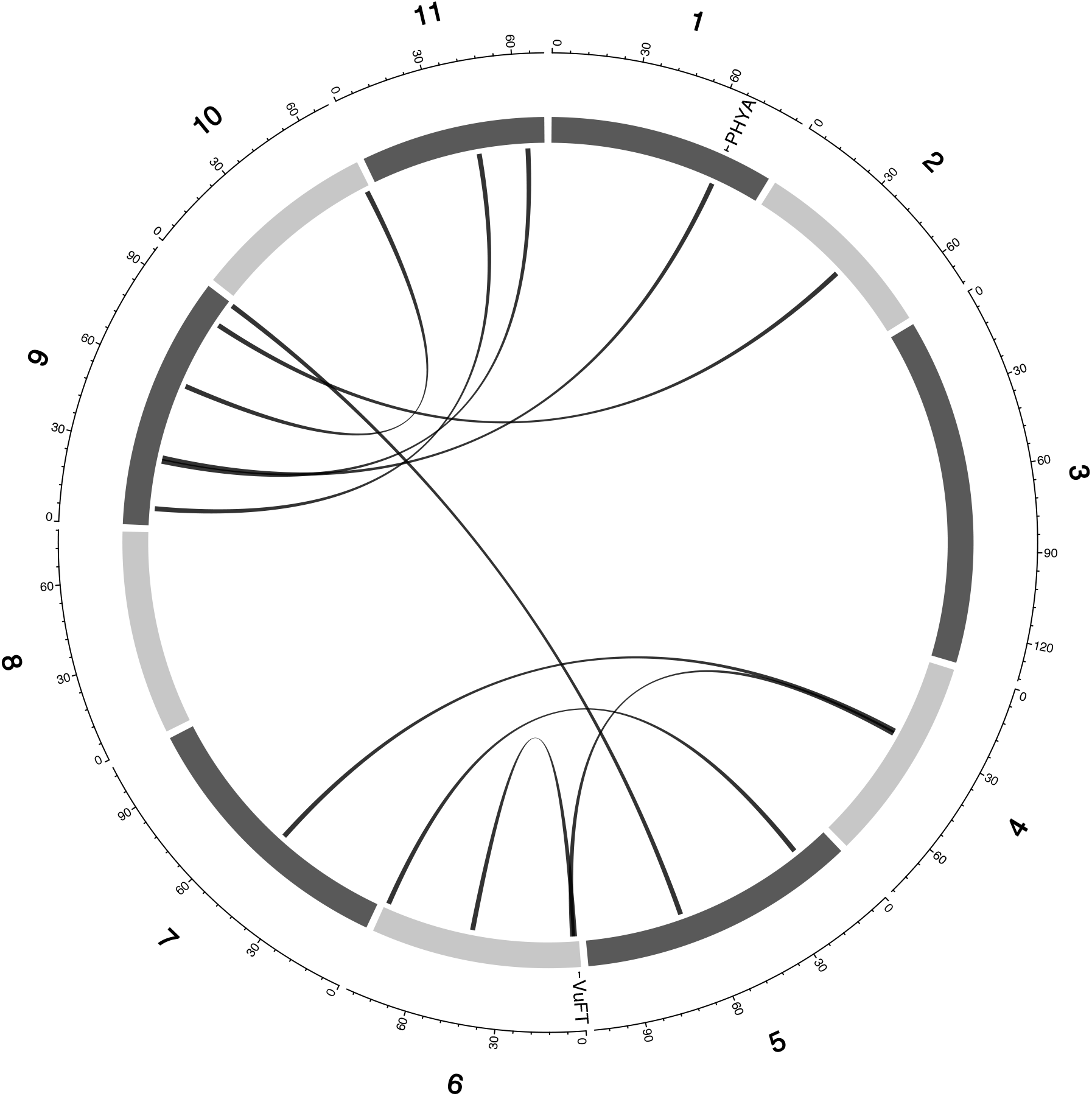
Epistatic QTL for FLT_BLUP for MAGIC population. Chromosomes are shown in shades of gray, two-way interacting loci are connected with black solid lines, and colocalized a priori genes are texts between chromosomes and genetic map.

**Figure 4:**
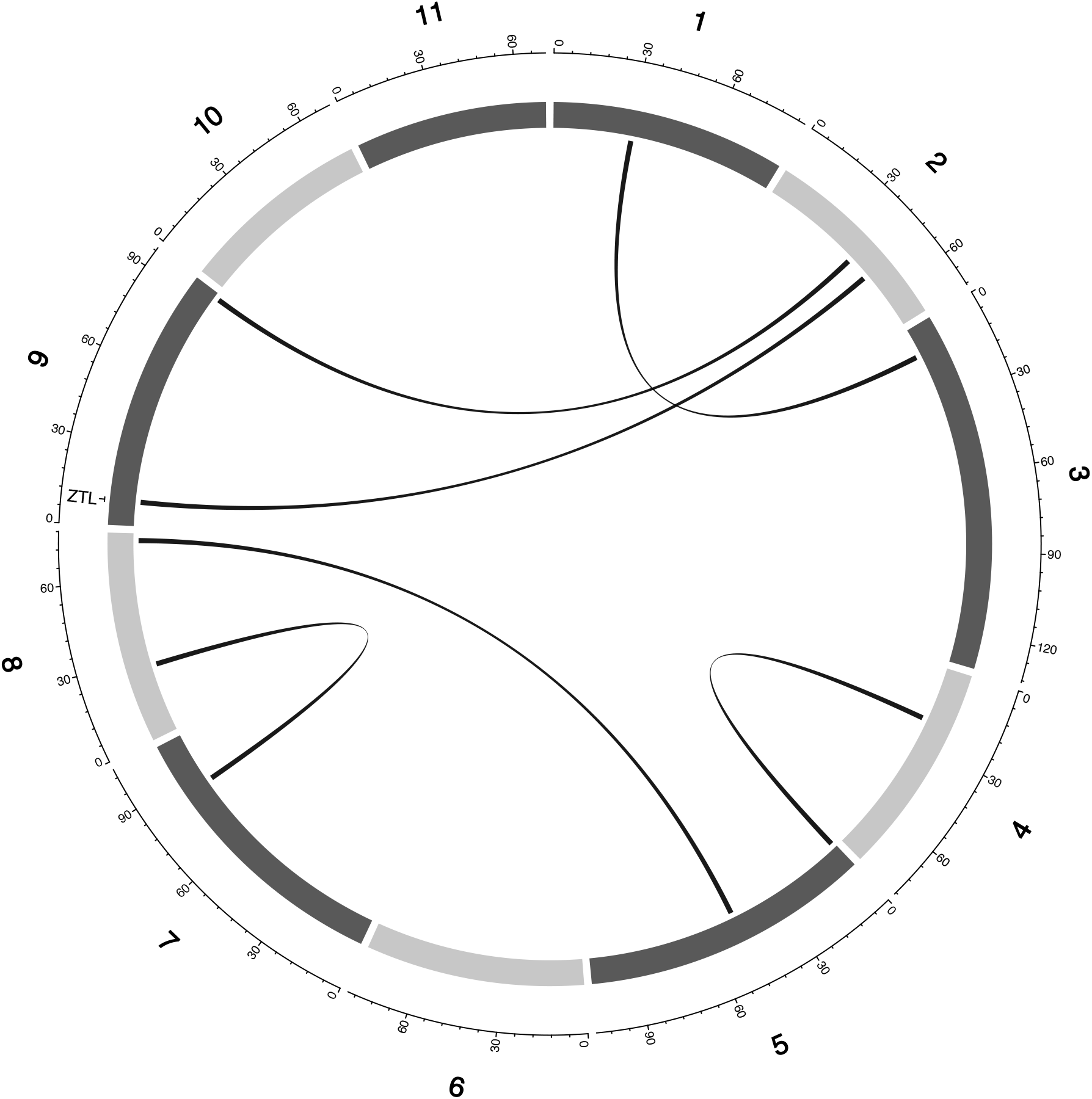
Epistatic QTL for MAT_BLUP in MAGIC population. Chromosomes are shown in shades of gray, two-way interacting loci are connected with black solid lines, and colocalized a priori genes are texts between chromosomes and genetic map.

**Figure 5:**
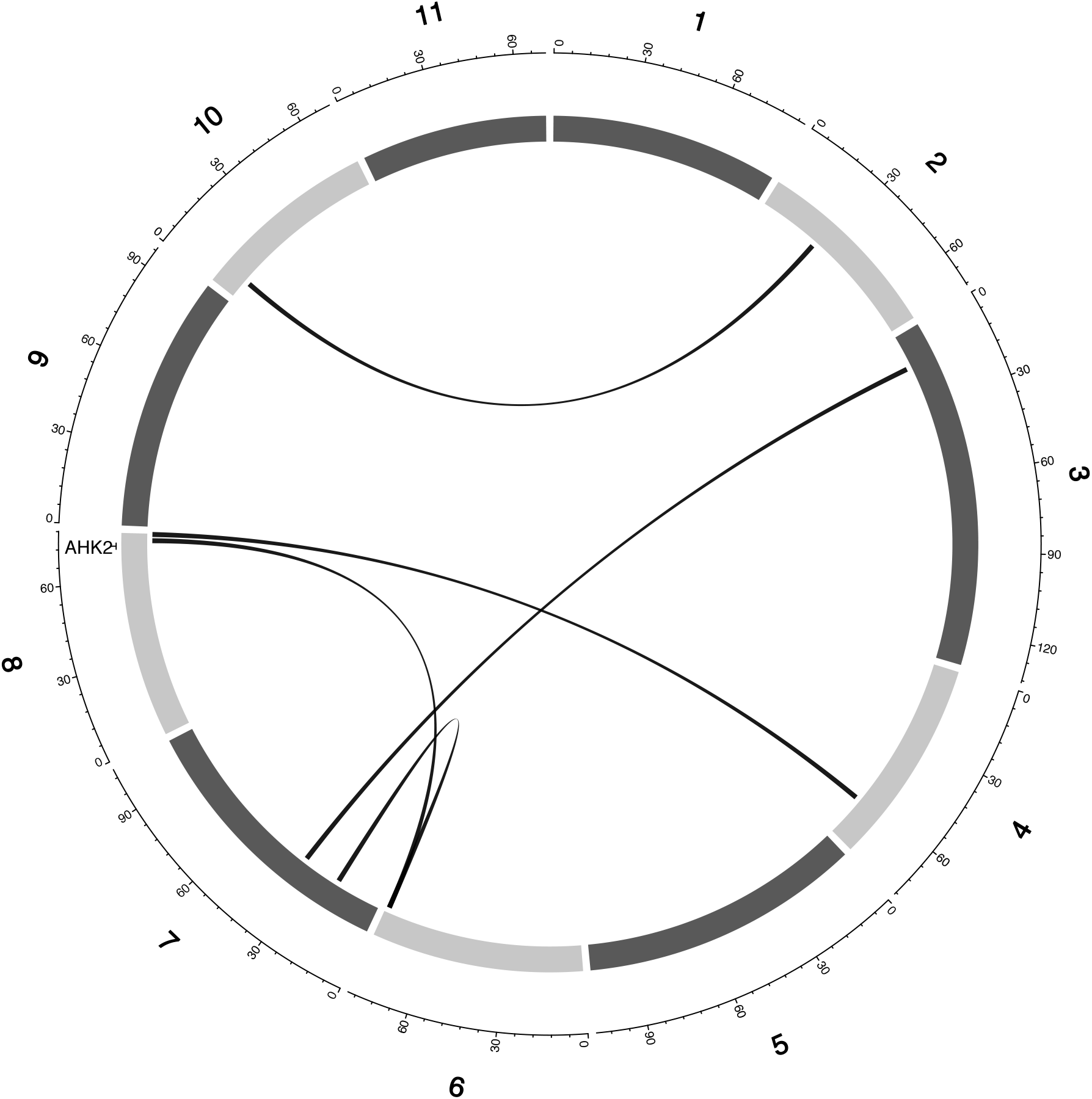
Epistatic QTL for MAT_BLUP in MAGIC population. Chromosomes are shown in shades of gray, two-way interacting loci are connected with black solid lines, and colocalized a priori genes are texts between chromosomes and genetic map.

Flowering time is an important adaptive trait in breeding. In this study, our results demonstrated that the flowering time variation in cowpea is due to large and moderate main effects and epistatic loci (Table S3 and Table S4). Epistatic loci underlie flowering time in both selfing (Huang *et al*., 2013; Juenger *et al*., 2005; Komeda, 2004;Mathew *et al*., 2018) (Chen *et al*., 2018)(Li *et al*., 2018) and outcrossing (Buckler *et al*., 2009; Durand *et al*., 2012) species. In addition, the effect size of flowering time loci differs between selfing and out crossing species as QTL effect sizes are large in the former (Lin, Schertz and Paterson, 1995; Maurer *et al*., 2015) and small in the later (Buckler *et al*., 2009). In the present study, the large effects (up to 25% PVE and additive effect of 7 days) flowering time loci were only identified under long day photoperiod and not under short-day photoperiod (Table S3 and Table S4). The loci detected under short day photoperiod were of moderate effects (PVE=1%-10% and maximum additive effect size of 2 days). A trait’s genetic architecture is a reflection of its stability over evolutionary time and traits subjected to strong recent selection were characterized with large effect loci (Brown *et al*., 2011). Our result suggests that cowpea flowering time adaptation to long-day photoperiod has undergone a recent selection compared to flowering time under short-day photoperiod.

### Distinct and common genetic regulators underlie flowering time

Conserved genetic pathways often underlie traits in plant species (Liu *et al*., 2013; Huang *et al*., 2017). Examination of colocalizations between a priori genes and main effect and epistatic QTL in this study identified putative cowpea orthologs of *A. thaliana* and *G. max* flowering time and seed size genes that may be underlie phenotypic variation in cowpea. Flowering time is affected by photoperiodicity and regulated by a network of genes (Sasaki, Frommlet and Nordborg, 2017) involved in floral initiation, circadian clock regulation, and photoreception (Lin, 2002). Photoperiod impacted days to flowering time as observed from the norm of reaction plot for cowpea MAGIC flowering time data which showed drastic reductions in days to flowering for RILs under short day compared to long days (Figure 1). Our mapping results (main effect and epistatic) showed both unique and common loci underlying flowering time under both long and short photoperiod (Figure 1; Figure S4–S8). In addition, certain *a priori genes* were unique to either flowering time under long day or short day. For instance, cowpea putative orthologs of photoreceptors (*PHY A* [Vigun01g205500] and *PHY E* [Vigun09g050600]) and circadian clock gene *PHYTOCLOCK1* (*PCL1* [Vigun11g157600]) colocalized with only QTL associated with flowering time under long day, while cowpea putative orthologs of circadian clock genes (*Time for Coffee* [*TIC* (Vigun07g241700)] and *Zeitlupe* [ZTL]) colocalized with only QTL associated with flowering time under short day. However, the cowpea putative ortholog of photoperiod gene *CONSTANS* (*CO* [Vigun05g024400]) colocalized with QTL associated with flowering time under both long and short days. Thus, our study suggests that distinct and common genetic regulators control flowering time adaptation to both long and short-day photoperiod in cowpea. Further studies utilizing functional approaches will be helpful to decipher gene regulation patterns under both long and short photoperiod in cowpea.

### Genetic architecture influenced GS and MAS performance

GS models differ in their efficiency to capture complex cryptic interactions among genetic markers (de Oliveira Couto *et al*., 2017). The traits evaluated in this study are controlled by both main effect and epistatic loci. In this study, comparison among the GS models showed that parametric and semi-parametric GS models outperformed non-parametric GS model for all traits. SVR, a non-parametric model had the least prediction accuracy and coincidence index and also had the highest bias (Figure S12 and S13). Previous studies have shown that semi-parametric and non-parametric models increased prediction accuracy under epistatic genetic architecture (Howard, Carriquiry and Beavis, 2014; Jacquin, Cao and Ahmadi, 2016). In this study, none of semi-parametric and non-parametric models outperformed parametric models (Figure 6 and 7). Some of the studies comparing the performance of parametric, semi-parametric and non-parametric GS models were based on simulations of traits controlled solely by epistatic genetic architectures. Therefore, the performance of the models under simulated combined genetic effects (additive + epistasis) is not well understood. The comparable performance of RKHS to RRBLUP (parametric model) in this study in terms of prediction accuracy, coincidence index, and bias estimates, attests to RKHS ability to capture both additive and epistatic interactions (Gianola, Fernando and Stella, 2006; Gianola and Van Kaam, 2008; De Los Campos *et al*., 2010; Gota and Gianola, 2014) for both prediction accuracy and selection of top performing lines. The performance of GS models’ is often indistinguishable and RRBLUP has been recommended as an efficient parametric GS model (Heslot *et al*., 2012; Lipka *et al*., 2015). SVR had the worst performance with extremely high bias estimates.

**Figure 6:**
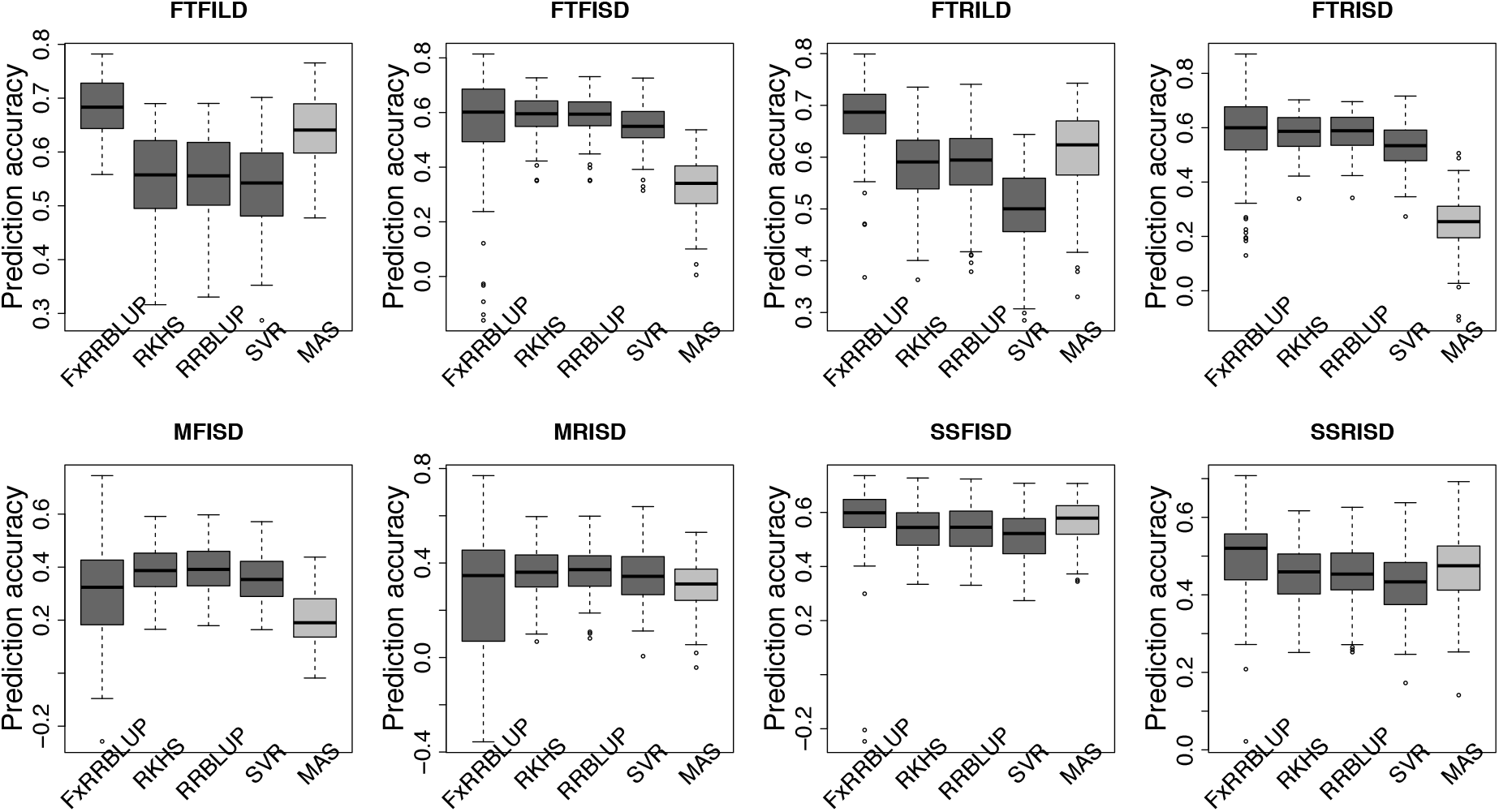
Comparison of prediction accuracy across GS and MAS models. Boxplots in each panel showed the distribution of prediction accuracy values across 100 cycles for FxRRBLUP (Ridge Regression Best Linear Unbiased Prediction: Parametric model with fixed effects), RKHS (Reproducing Kernel Hilbert Space; Semi-Parametric model), RRBLUP (Ridge Regression Best Linear Unbiased Prediction: Parametric model with no fixed effects), SVR (Support Vector Regression: Non-Parametric model), and MAS (Marker Assisted Selection) for flowering time under full irrigation and long day (FTFILD), flowering time under restricted irrigation and long day (FTRILD), flowering time under full irrigation and short day (FTFISD), flowering time under restricted irrigation and short day (FTRISD), maturity under full irrigation and short day (MFISD), maturity under restricted irrigation and short day, seed size under full irrigation and short day (SSFISD), and seed size under restricted irrigation and short day (SSRISD).

**Figure 7:**
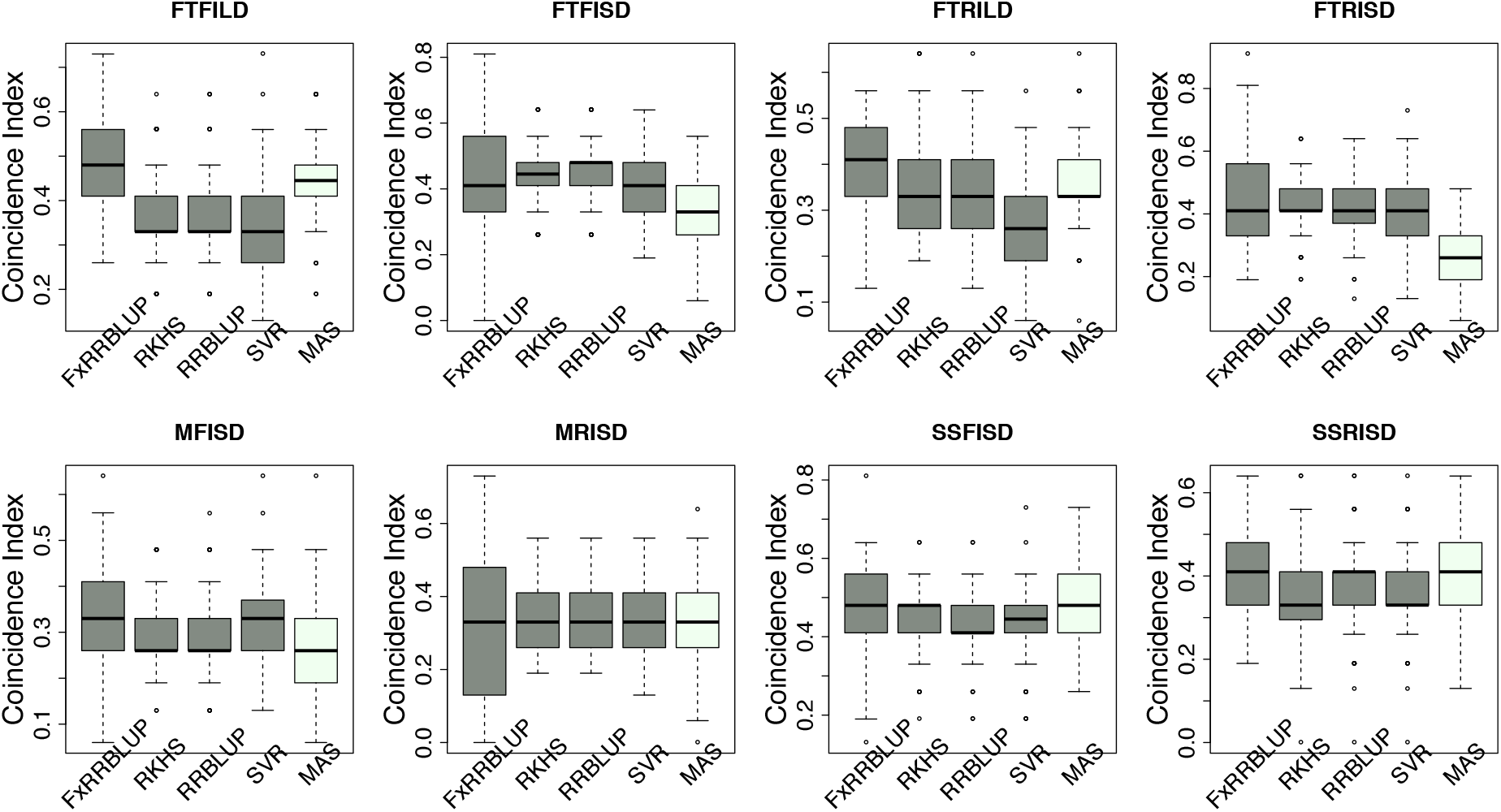
Comparison of coincidence index across GS and MAS models. Boxplots in each panel showed the distribution of coincidence index values across 100 cycles for FxRRBLUP (Ridge Regression Best Linear Unbiased Prediction: Parametric model with fixed effects), RKHS (Reproducing Kernel Hilbert Space; Semi-Parametric model), RRBLUP (Ridge Regression Best Linear Unbiased Prediction: Parametric model with no fixed effects), SVR (Support Vector Regression: Non-Parametric model), and MAS (Marker Assisted Selection) for flowering time under full irrigation and long day (FTFILD), flowering time under restricted irrigation and long day (FTRILD), flowering time under full irrigation and short day (FTFISD), flowering time under restricted irrigation and short day (FTRISD), maturity under full irrigation and short day (MFISD), maturity under restricted irrigation and short day, seed size under full irrigation and short day (SSFISD), and seed size under restricted irrigation and short day (SSRISD).

Understanding the genetic architecture of agronomic traits can help improve predictions (Hayes *et al*., 2010; Swami, 2010). Our study demonstrated that the effect size of QTL associated with a trait played a role in the performance of GS and MAS models. For instance, for traits controlled by both large and moderate effects loci (FTFILD, FTRILD, SSFISD, and SSRISD) parametric model with known loci as fixed effect (FxRRBLUP) followed by MAS outperformed the rest of the GS models (RRBLUP, RKHS, and SVR). The use of known QTL as fixed effects has been shown to increase prediction accuracy (Bernardo, 2014; Spindel *et al*., 2016) in parametric GS models. For traits that were controlled by moderate effects loci (FTFISD, FTRISD, MFISD, and MTRISD), our results showed that the two parametric GS models (FxRRBLUP and RRBLUP) and semi-parametric (RKHS) had similar prediction accuracy, however FxRRBLUP had higher bias than RRBLUP and RKHS (Figure S12 – S13). Accuracy of prediction is influenced by genetic architecture (Hayes *et al*., 2010). Furthermore, the performance of MAS in comparison to GS models in this study showed that large effects loci are important influencers of MAS (Bernardo, 2008). For small breeding programs in developing countries, MAS might be a prudent choice over GS for traits controlled only loci of large effects in cowpea since GS will require genotyping of more markers than MAS. The large effect QTL identified in this study can be transferred to different breeding populations because they were identified in a MAGIC population with wide genetic background (Chen *et al*., 2018; Huynh *et al*., 2018). Our study thus demonstrates that prior knowledge of the genetic architecture of a trait can help make informed decision about the best GEB method to employ in breeding.

### Experimental design considerations for GS in cowpea

An important consideration in this study is to provide recommendations to breeders on resources needed for the implementation of GS in cowpea. First, this study demonstrated that genomic prediction within the same photoperiod is more efficient than across different photoperiod (Figure S14). Prediction between irrigation regimes had similar performance. The differences observed for GS between photoperiods showed that genotype x environment (GxE) interaction is an important factor to consider in cowpea flowering time GS. Increased genetic gains were observed in GS approaches that modeled GxE interactions (Lopez-Cruz *et al*., 2015; Crossa *et al*., 2016; de Oliveira Couto *et al*., 2017). Second, our results showed that the size of the training population had an effect on prediction accuracy as prediction accuracy increased with increase in training population size. The size of a training population is an important factor influencing prediction accuracy (Liu *et al*., 2018) and studies have shown increase in prediction accuracy with increase in training population size in several crop species (Albrecht *et al*., 2011; Spindel *et al*., 2015). Third, increase in marker density only significantly increased prediction between 20-60% for FLT_BLUP and 40-60% and 60-80% for SS_BLUP (Figure S19). Though these differences were significant, the mean prediction accuracy values were close to each other for all marker densities (Figure S19A). If using 20% of markers (6424 SNPs) gave similar prediction accuracy as 32,130 SNPs; then it might be more cost efficient for a breeder to use a small marker density. For instance, for flowering time, 6424 SNPs gave a mean prediction accuracy of 0.665 and 32130 SNPs gave a prediction accuracy of 0.671, then it might be logical and cost efficient to use ~6000 markers for GS.

In summary, to the best of our knowledge, this is the first study that will characterize epistasis and provide insights into the underpinnings of genomic selection *versus* marker assisted selection in cowpea. Our study identified both main QTL and two-way epistatic loci underlying flowering time, maturity, and seed size. We also found that flowering time is under the control of both large and moderate effect loci similar to findings in other inbreeding species. The large effect QTL and their colocalized *a priori* genes identified in this study will serve as pedestal for future studies aimed at the molecular characterization of the genes underlying flowering time and seed size in cowpea. We demonstrated that prior knowledge of the genetic architecture of a trait can help make informed decision in GEB. Together, our findings in this study are relevant for crop improvement in both developed and developing countries.

## Acknowledgement

We express our gratitude to Prof. Timothy Close, Prof. Philip Roberts, Dr. Bao-Lam Huynh and their team at the University of California - Riverside, USA for their incredible contributions to cowpea genomics and the privilege to use the cowpea MAGIC population data for this study. The MAGIC population development, phenotyping, and genotyping was supported in large part by grants from the Generation Challenge Program of the Consultative Group on International Agricultural Research, with additional support from the USAID Feed the Future Innovation Lab for Collaborative Research on Grain Legumes (Cooperative Agreement EDH-A-00-07-00005), the USAID Feed the Future Innovation Lab for Climate Resilient Cowpea (Cooperative Agreement AID-OAA-A-13-00070), and NSF-BREAD (Advancing the Cowpea Genome for Food Security). We also thank Dr. Sandeep Marla, and Fanna Maina for helping with the manuscript review.

## Authors’ contributions

M.O.O. obtained data from UCR; concept by M.O.O and Z.H; M.O.O. and Z.H. analyzed the data; M.O.O, Z.H, and P.O.A wrote the manuscript. All authors read and approved the manuscript.

## Supporting information

All the R scripts used for analyses in the study are available at: https://github.com/marcbios/Cowpea.git

## Tables

### Supplementary figures

**Figure S 1:**
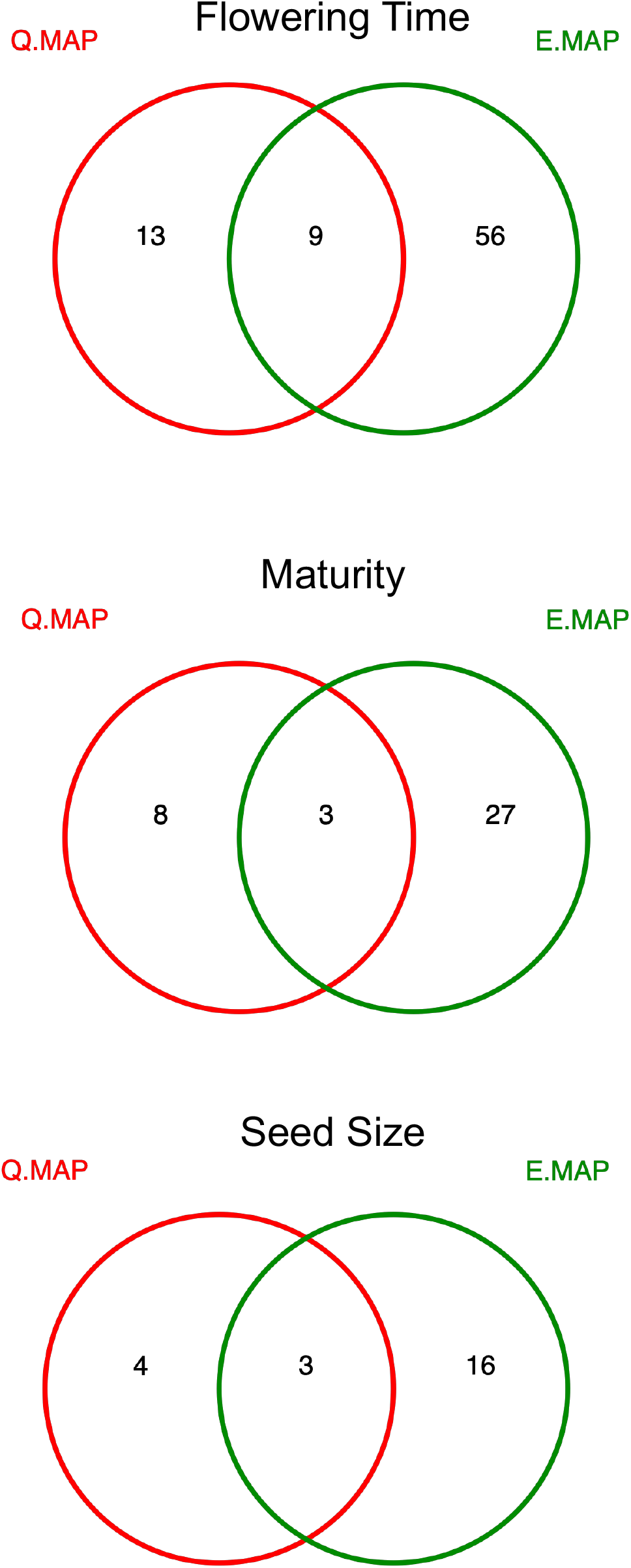
Venn diagram of QTL overlap between main effect QTL mapping (Q.MAP) and epistasis mapping (E.MAP) for flowering time, maturity, and seed size.

**Figure S 2:**
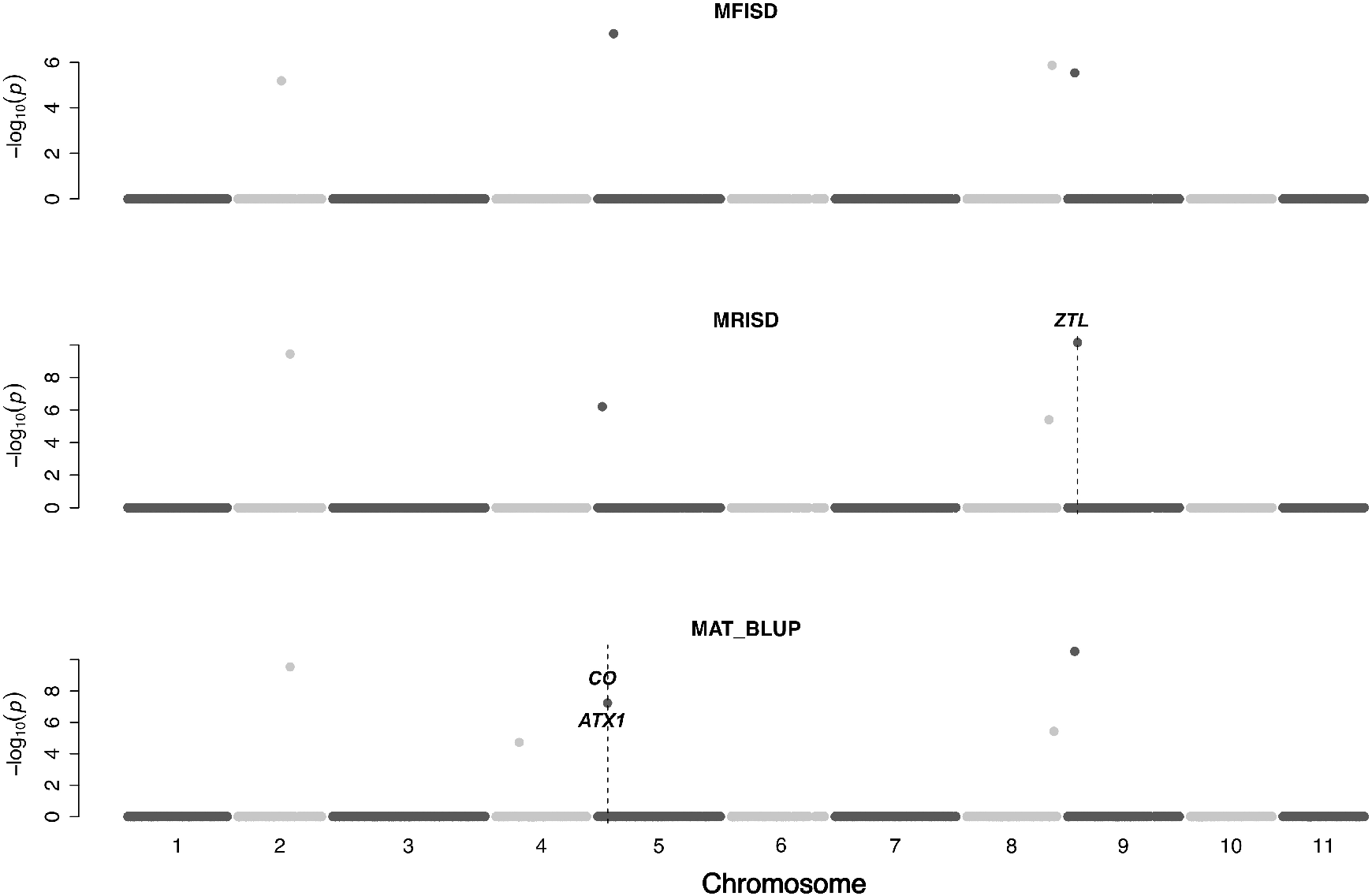
QTL plot for maturity traits in the cowpea MAGIC population. QTL plots for maturity under full irrigation and short day (MFISD), maturity under restricted irrigation and short day (MRISD), and BLUPs of environments (MAT_BLUP). The chromosome numbers are located on the x-axis and the negative log of the P-values on the y-axis. The genetic position of the colocalization between QTL and *a priori* genes are indicated by broken vertical lines. The texts displayed on the vertical broken lines are the names of *a priori genes*.

**Figure S 3:**
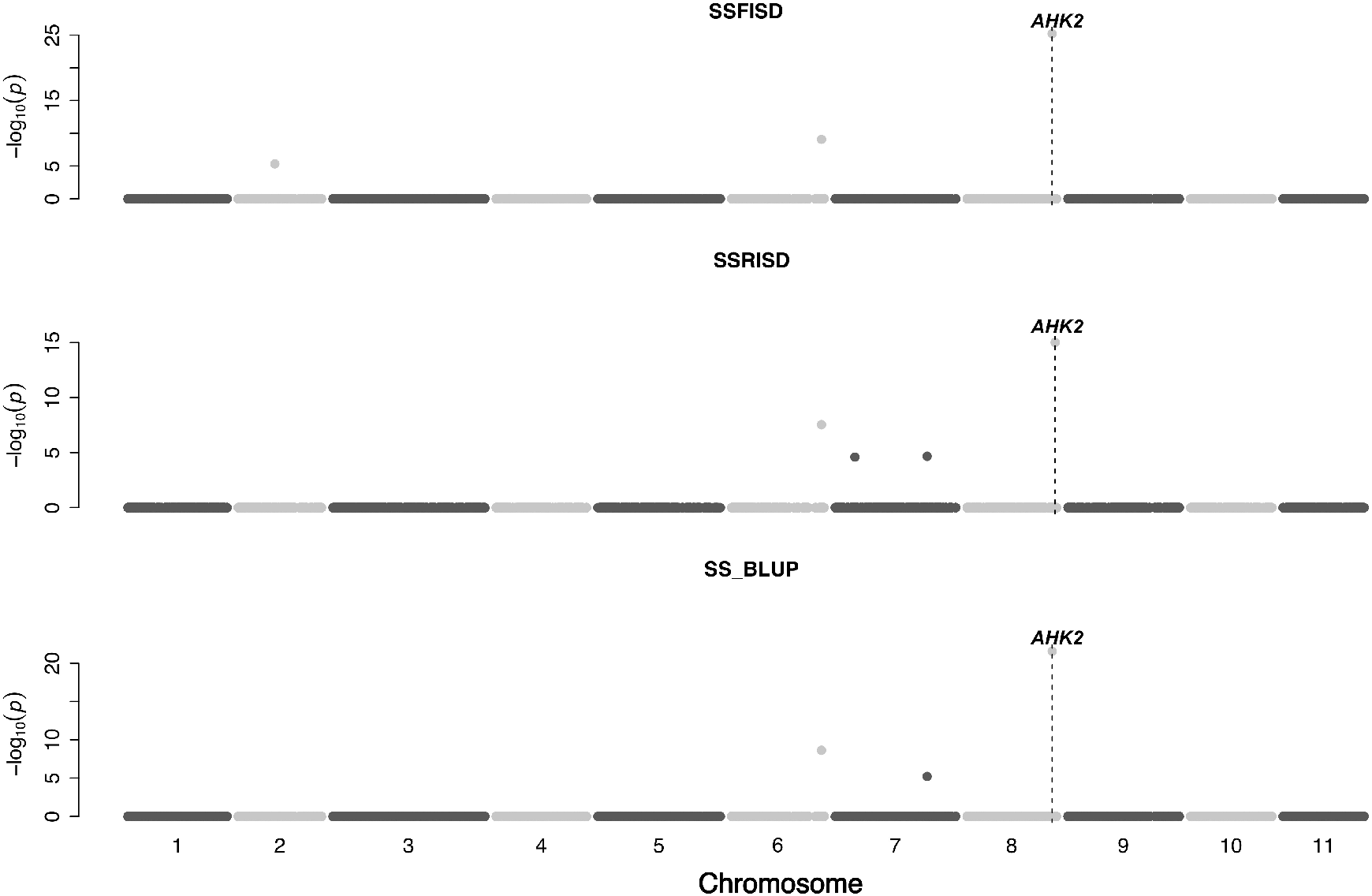
QTL plot for seed size traits in the cowpea MAGIC population. QTL plots for maturity under full irrigation and short day (SSFISD), maturity under restricted irrigation and short day (SSRISD), and BLUPs of environments (SS_BLUP). The chromosome numbers are located on the x-axis and the negative log of the *P*-values on the y-axis. The genetic position of the colocalization between QTL and *a priori* genes are indicated by broken vertical lines. The texts displayed on the vertical broken lines are the names of *a priori genes*.

**Figure S 4:**
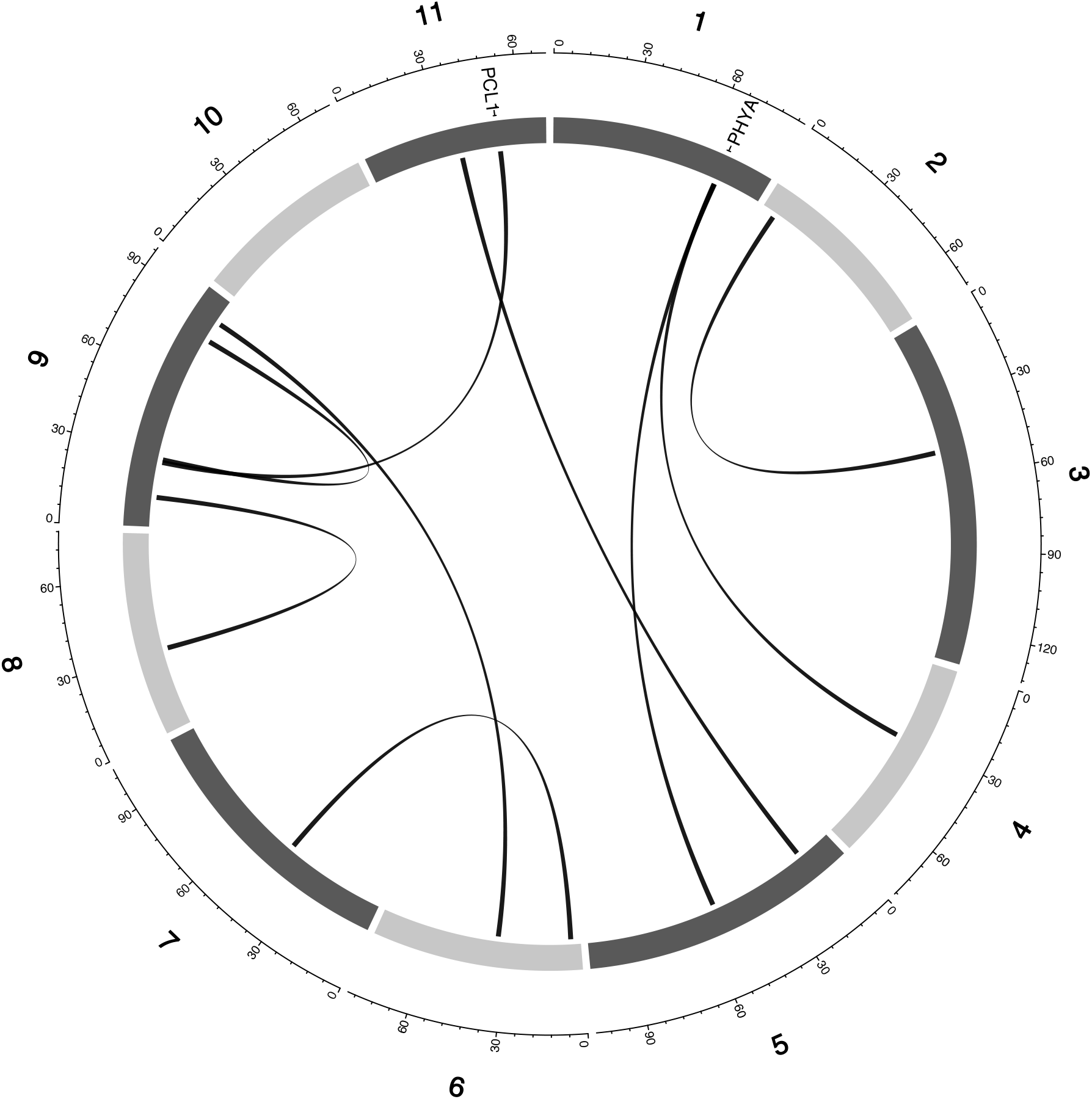
Genetic map of the cowpea multiparent advanced generation inter-cross population (MAGIC) with pairwise interactions between epistatic QTL for FTFILD (Flowering time under full irrigation and long day). Chromosomes are shown in shades of gray, two-way interacting loci are connected with black solid lines, and colocalized *a priori* genes are texts between chromosomes and genetic map.

**Figure S 5:**
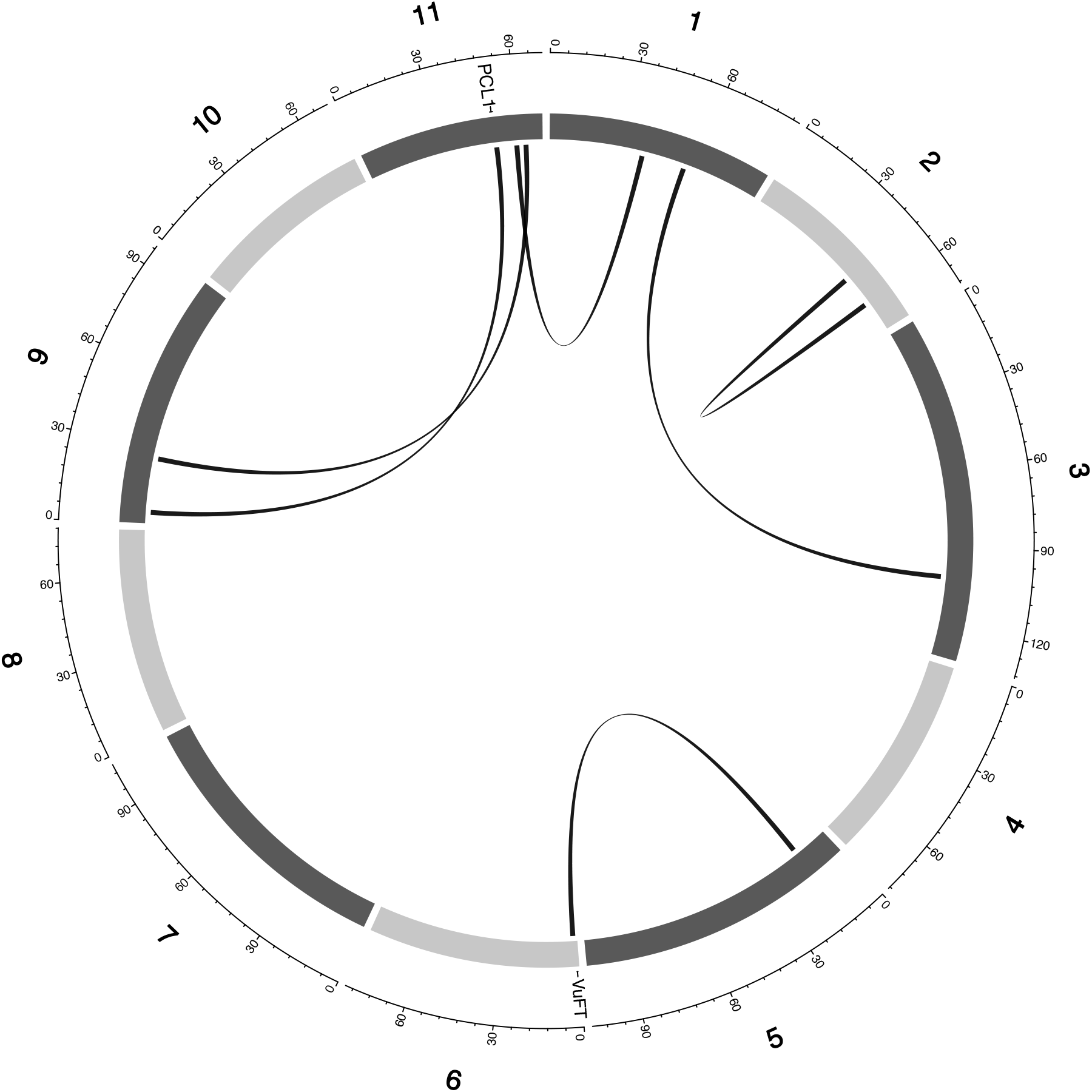
Genetic map of the cowpea multiparent advanced generation inter-cross population (MAGIC) with pairwise interactions between epistatic QTL for FTRILD (Flowering time under restricted irrigation and long day). Chromosomes are shown in shades of gray, two-way interacting loci are connected with black solid lines, and colocalized a priori genes are texts between chromosomes and genetic map.

**Figure S 6:**
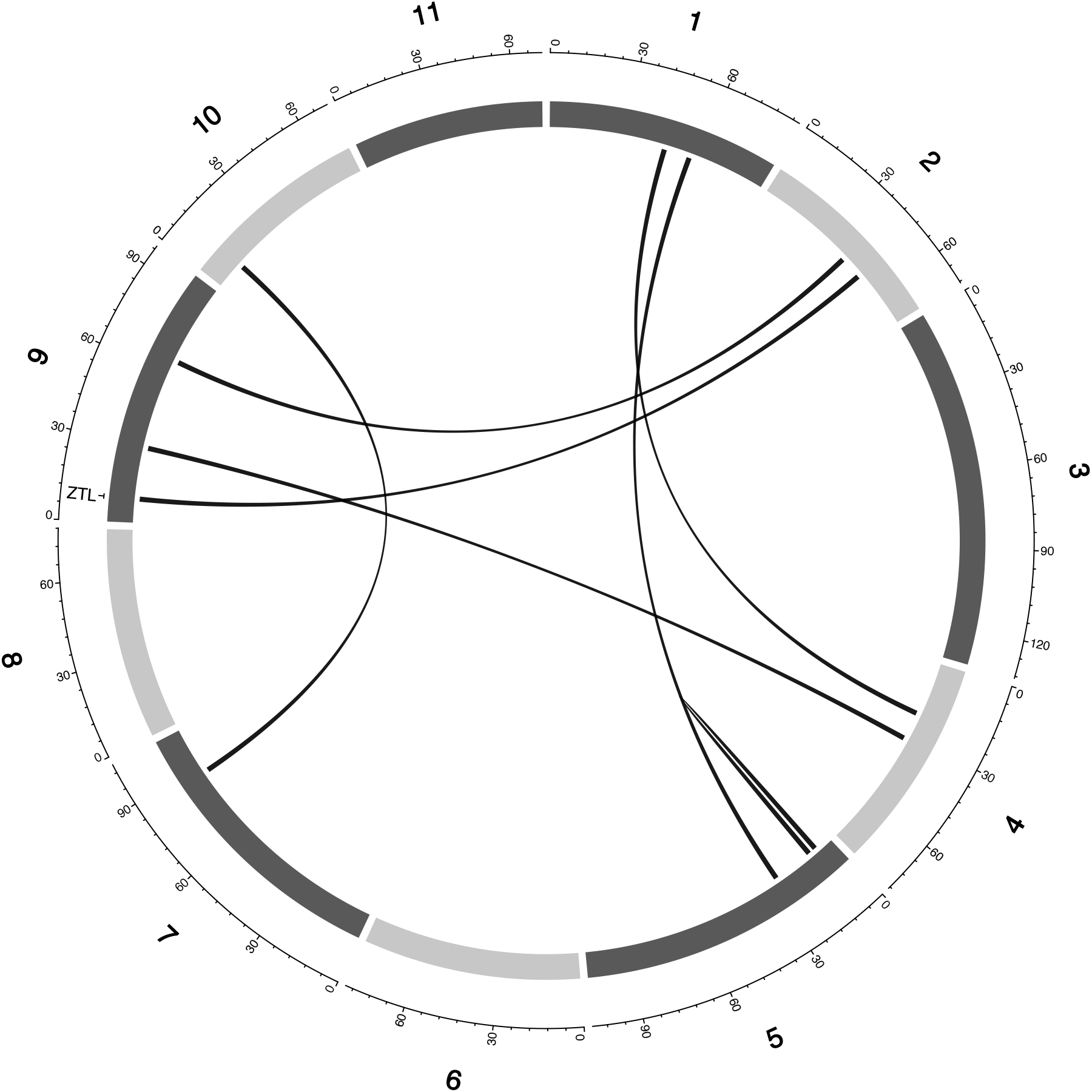
Genetic map of the cowpea multiparent advanced generation inter-cross population (MAGIC) with pairwise interactions between epistatic QTL for FTFISD (Flowering time under full irrigation and short day). Chromosomes are shown in shades of gray, two-way interacting loci are connected with black solid lines, and colocalized a priori genes are texts between chromosomes and genetic map.

**Figure S 7:**
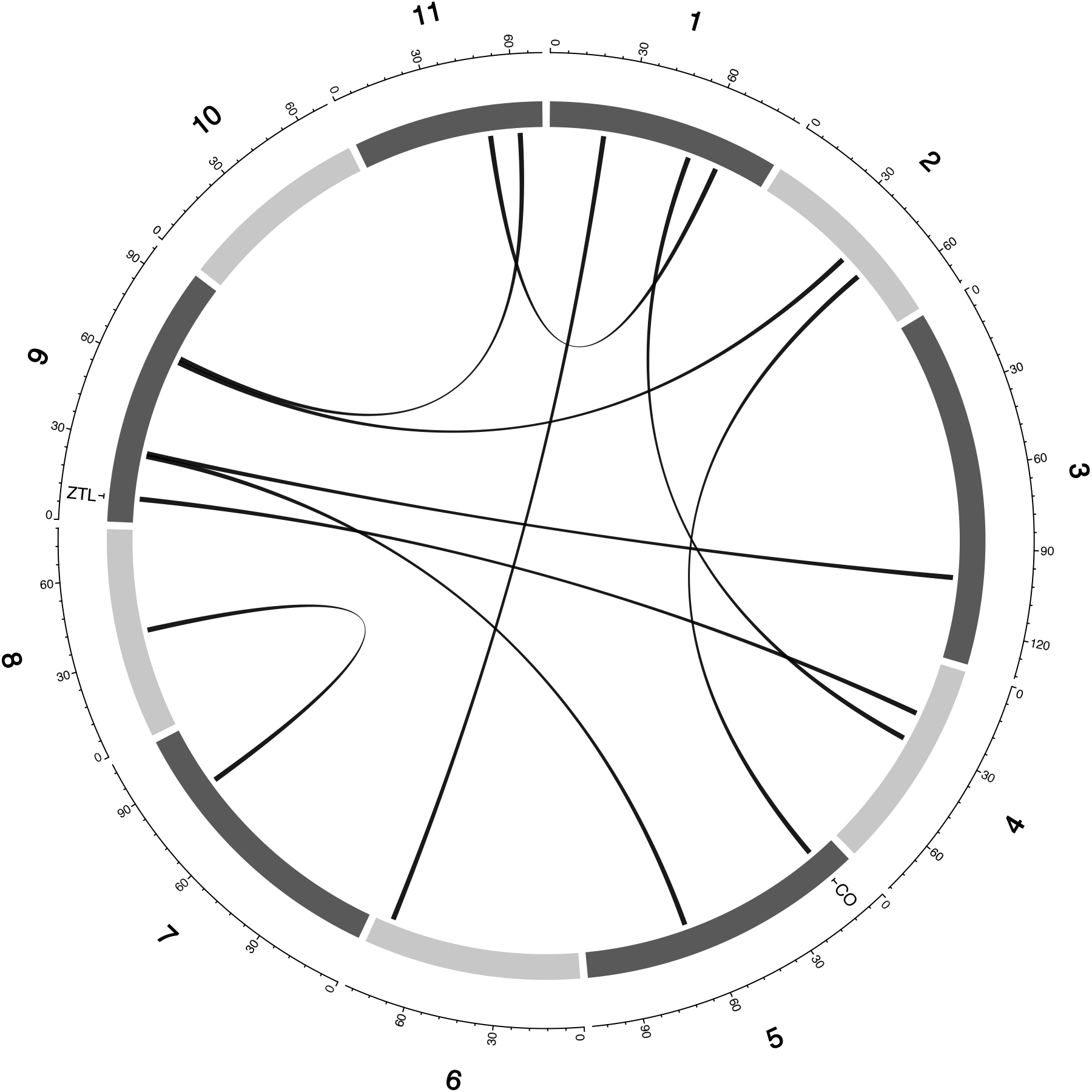
Genetic map of the cowpea multiparent advanced generation inter-cross population (MAGIC) with pairwise interactions between epistatic QTL for FTRISD (Flowering time under restricted irrigation and short day). Chromosomes are shown in shades of gray, two-way interacting loci are connected with black solid lines, and colocalized a priori genes are texts between chromosomes and genetic map.

**Figure S 8:**
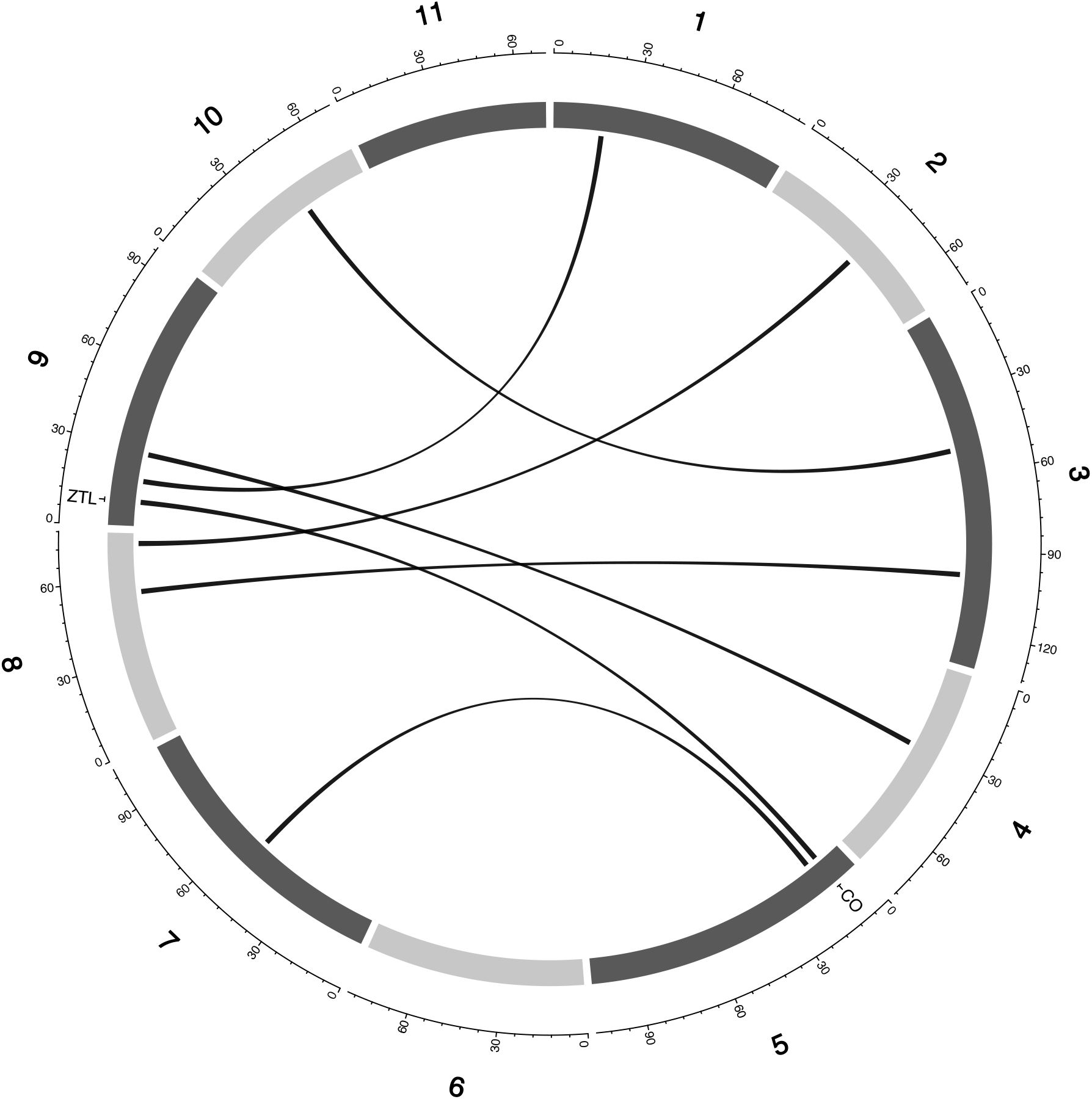
Genetic map of the cowpea multiparent advanced generation inter-cross population (MAGIC) with pairwise interactions between epistatic QTL for MFISD (Maturity under full irrigation and short day). Chromosomes are shown in shades of gray, two-way interacting loci are connected with black solid lines, and colocalized a priori genes are texts between chromosomes and genetic map.

**Figure S 9:**
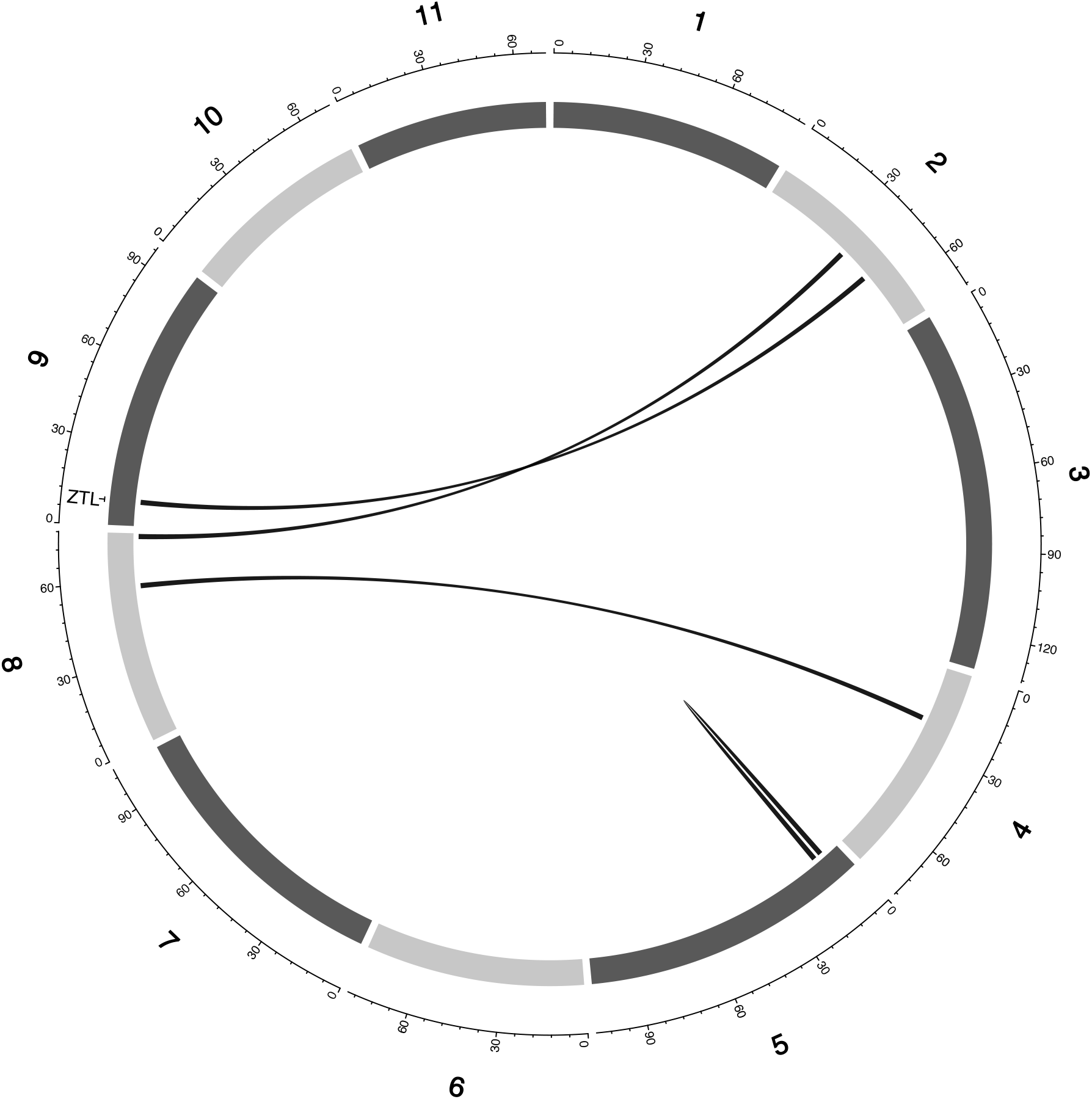
Genetic map of the cowpea multiparent advanced generation inter-cross population (MAGIC) with pairwise interactions between epistatic QTL for MRISD (Maturity under restricted irrigation and short day). Chromosomes are shown in shades of gray, two-way interacting loci are connected with black solid lines, and colocalized a priori genes are texts between chromosomes and genetic map.

**Figure S 10:**
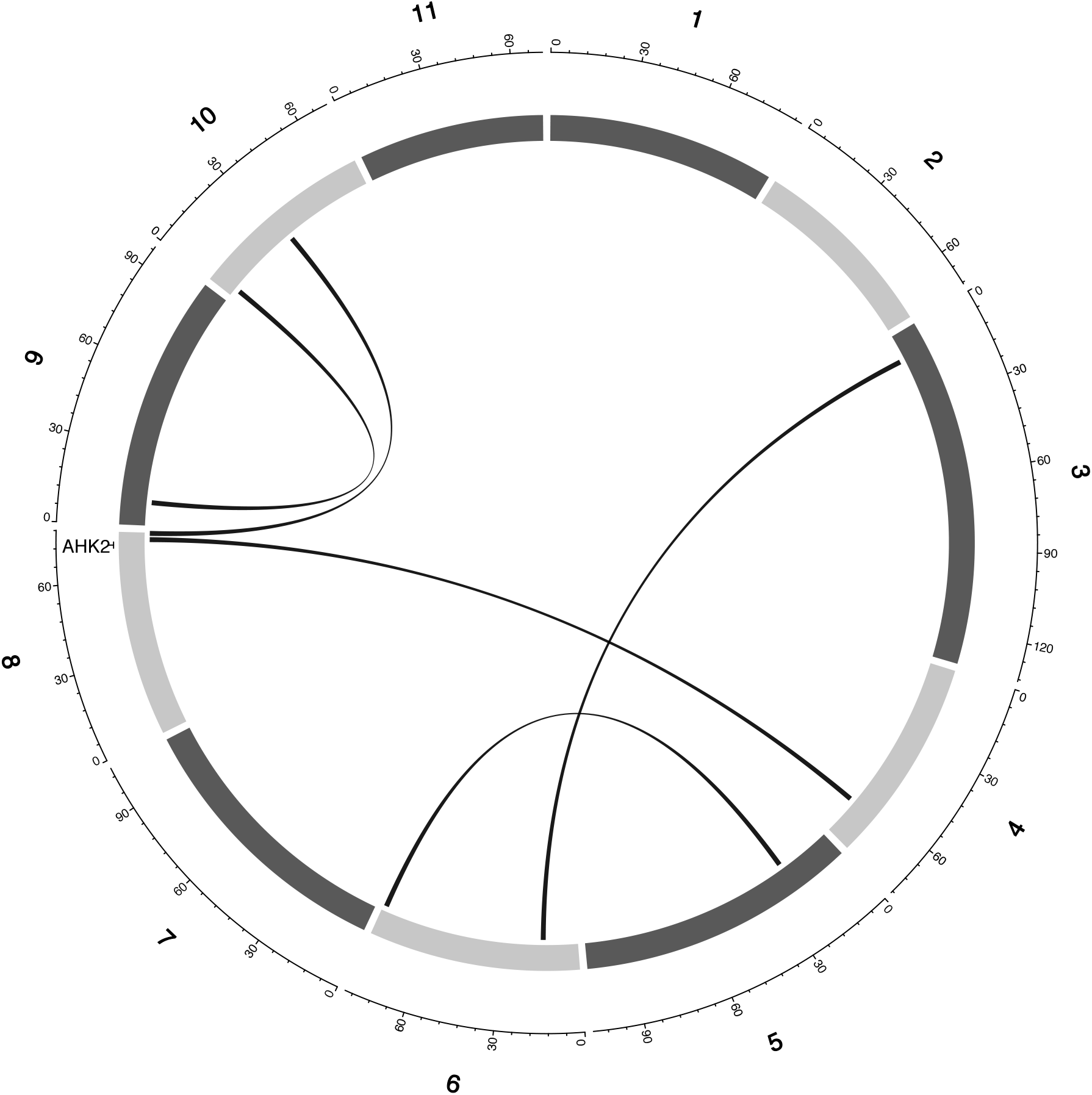
Genetic map of the cowpea multiparent advanced generation inter-cross population (MAGIC) with pairwise interactions between epistatic QTL for SSFISD (Seed Size under full irrigation and short day). Chromosomes are shown in shades of gray, two-way interacting loci are connected with black solid lines, and colocalized a priori genes are texts between chromosomes and genetic map.

**Figure S 11:**
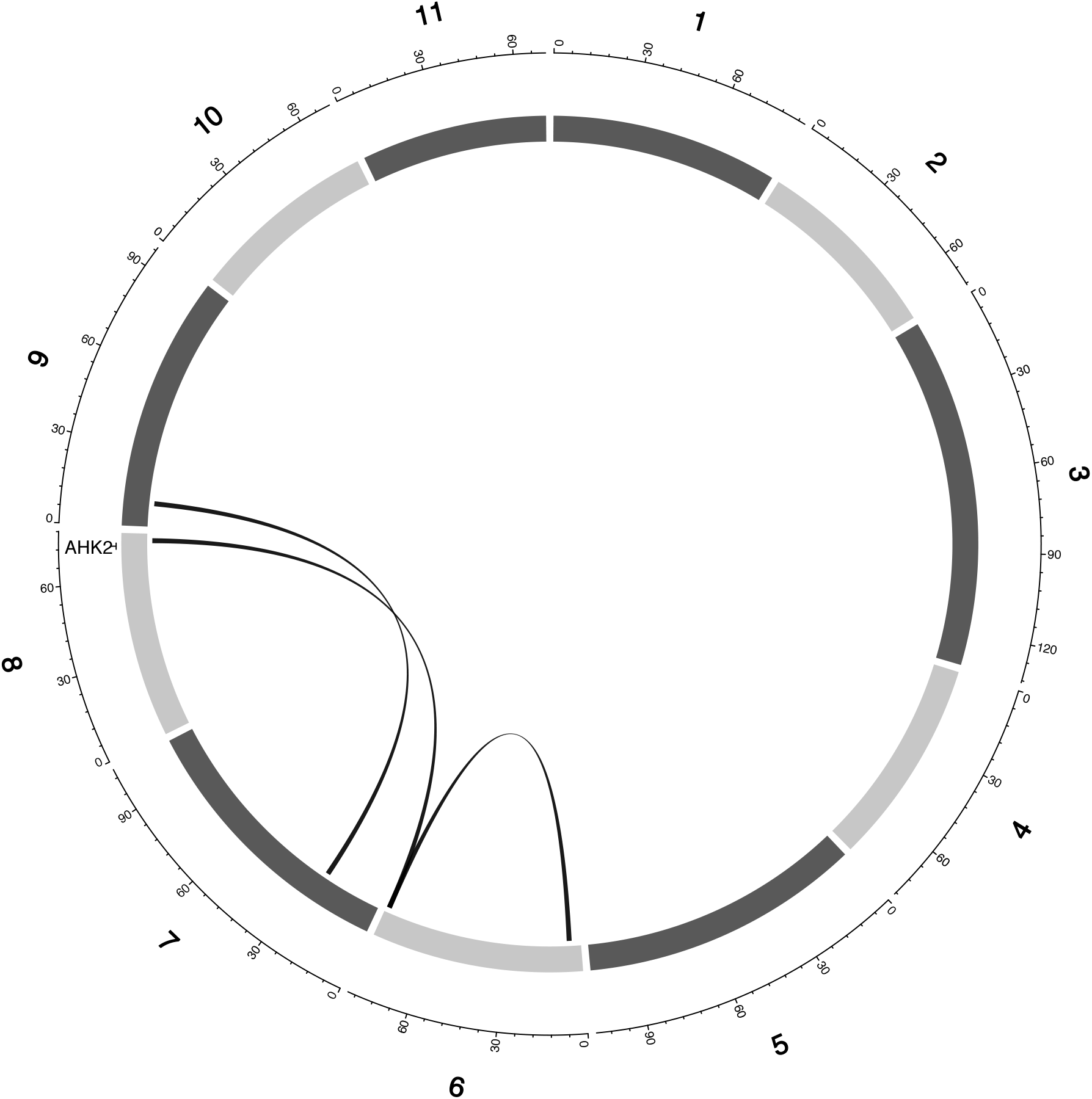
Genetic map of the cowpea multiparent advanced generation inter-cross population (MAGIC) with pairwise interactions between epistatic QTL for SSRISD (Seed size under restricted irrigation and short day). Chromosomes are shown in shades of gray, two-way interacting loci are connected with black solid lines, and colocalized a priori genes are texts between chromosomes and genetic map.

**Figure S 12:**
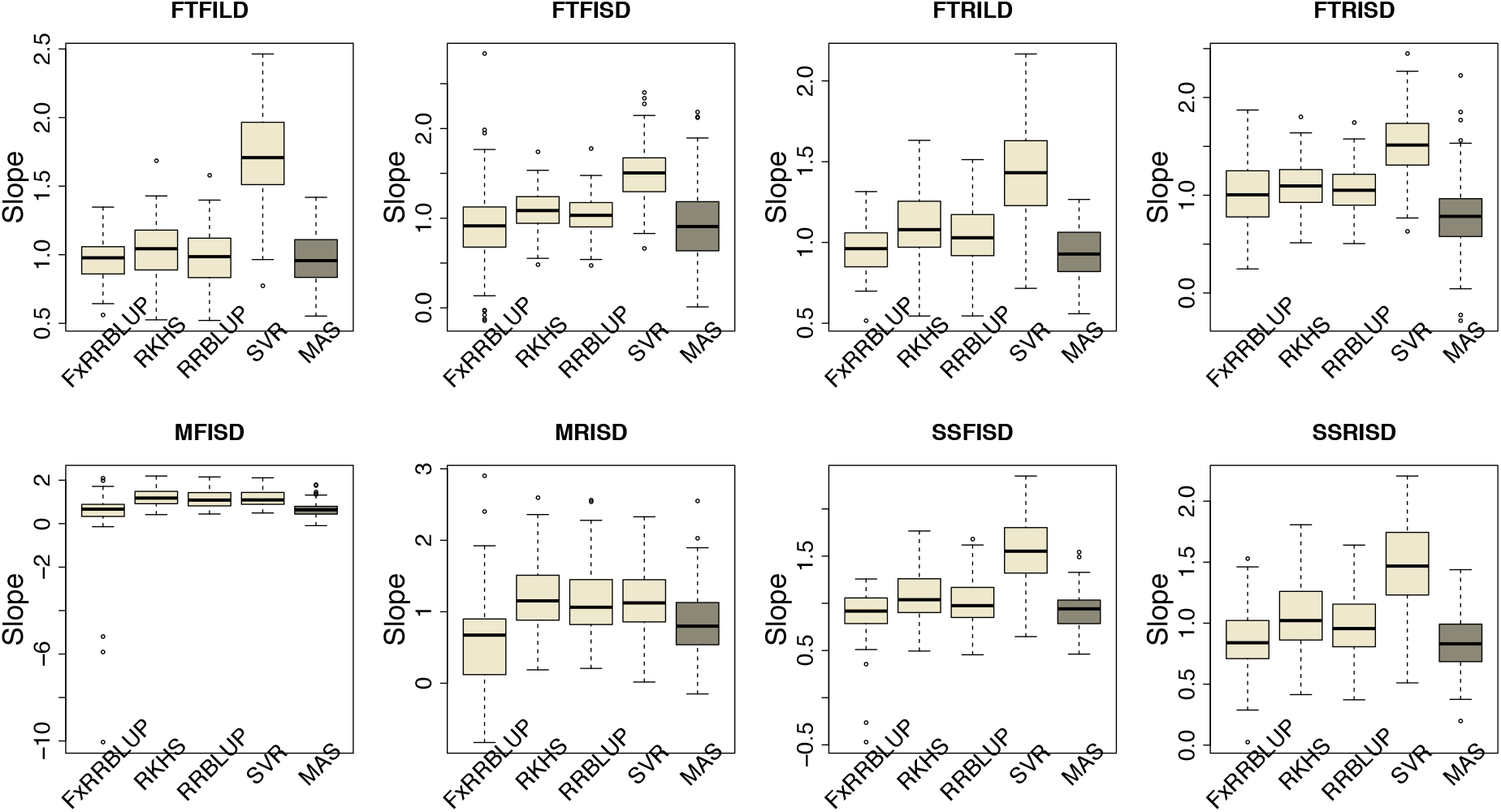
Comparison of Slope of regression between observed and predicted trait values across GS and MAS models. Boxplots in each panel showed the distribution of slope values across 100 cycles for FxRRBLUP (Ridge Regression Best Linear Unbiased Prediction: Parametric model with fixed effects), RKHS (Reproducing Kernel Hilbert Space; Semi-Parametric model), RRBLUP (Ridge Regression Best Linear Unbiased Prediction: Parametric model with no fixed effects), SVR (Support Vector Regression: Non-Parametric model), and MAS (Marker Assisted Selection) for flowering time under full irrigation and long day (FTFILD), flowering time under restricted irrigation and long day (FTRILD), flowering time under full irrigation and short day (FTFISD), flowering time under restricted irrigation and short day (FTRISD), maturity under full irrigation and short day (MFISD), maturity under restricted irrigation and short day, seed size under full irrigation and short day (SSFISD), and seed size under restricted irrigation and short day (SSRISD).

**Figure S 13:**
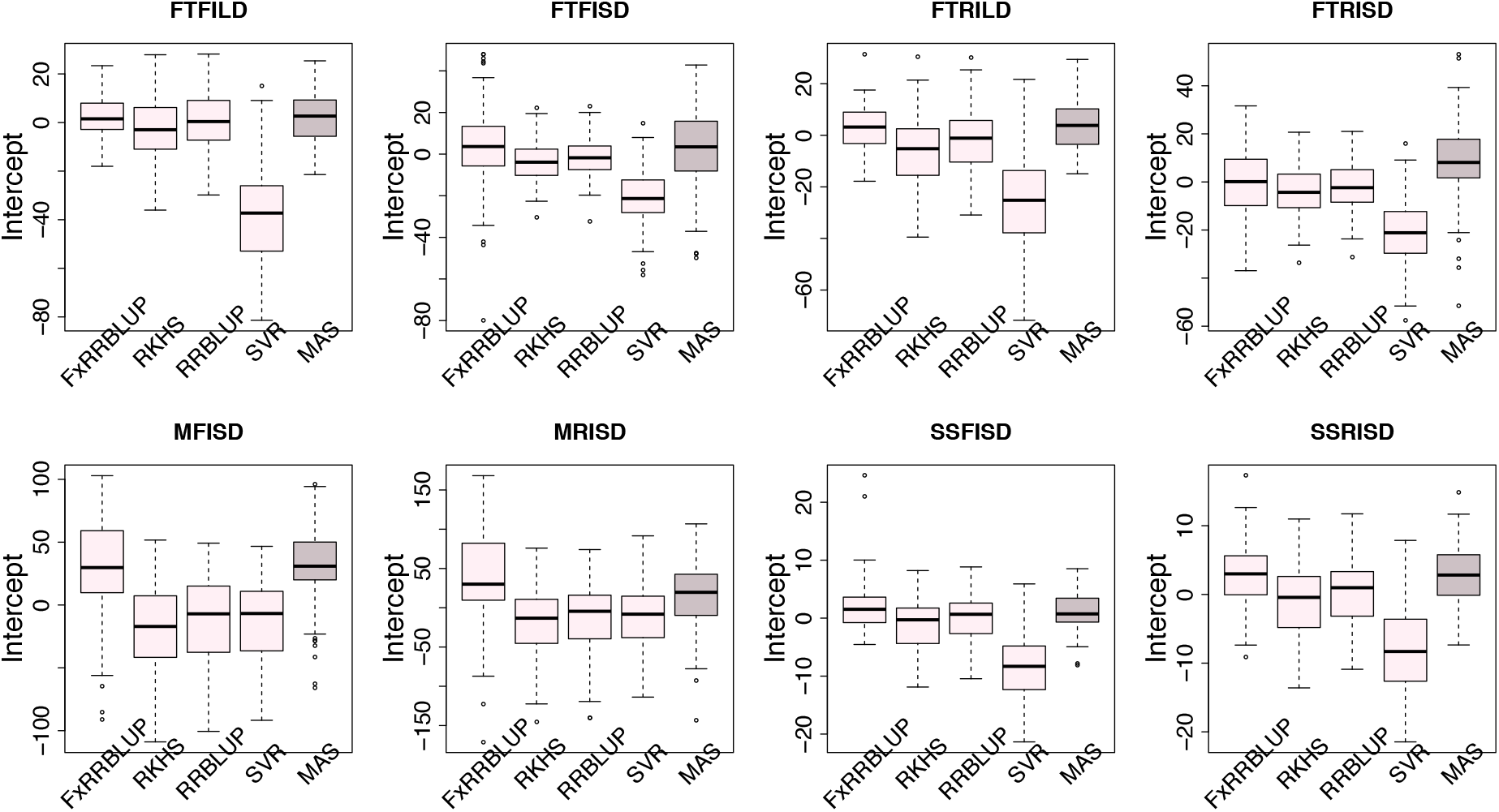
Comparison of intercept of regression between observed and predicted trait values across GS and MAS models. Boxplots in each panel showed the distribution of intercept values across 100 cycles for FxRRBLUP (Ridge Regression Best Linear Unbiased Prediction: Parametric model with fixed effects), RKHS (Reproducing Kernel Hilbert Space; Semi-Parametric model), RRBLUP (Ridge Regression Best Linear Unbiased Prediction: Parametric model with no fixed effects), SVR (Support Vector Regression: Non-Parametric model), and MAS (Marker Assisted Selection) for flowering time under full irrigation and long day (FTFILD), flowering time under restricted irrigation and long day (FTRILD), flowering time under full irrigation and short day (FTFISD), flowering time under restricted irrigation and short day (FTRISD), maturity under full irrigation and short day (MFISD), maturity under restricted irrigation and short day, seed size under full irrigation and short day (SSFISD), and seed size under restricted irrigation and short day (SSRISD).

**Figure S 14:**
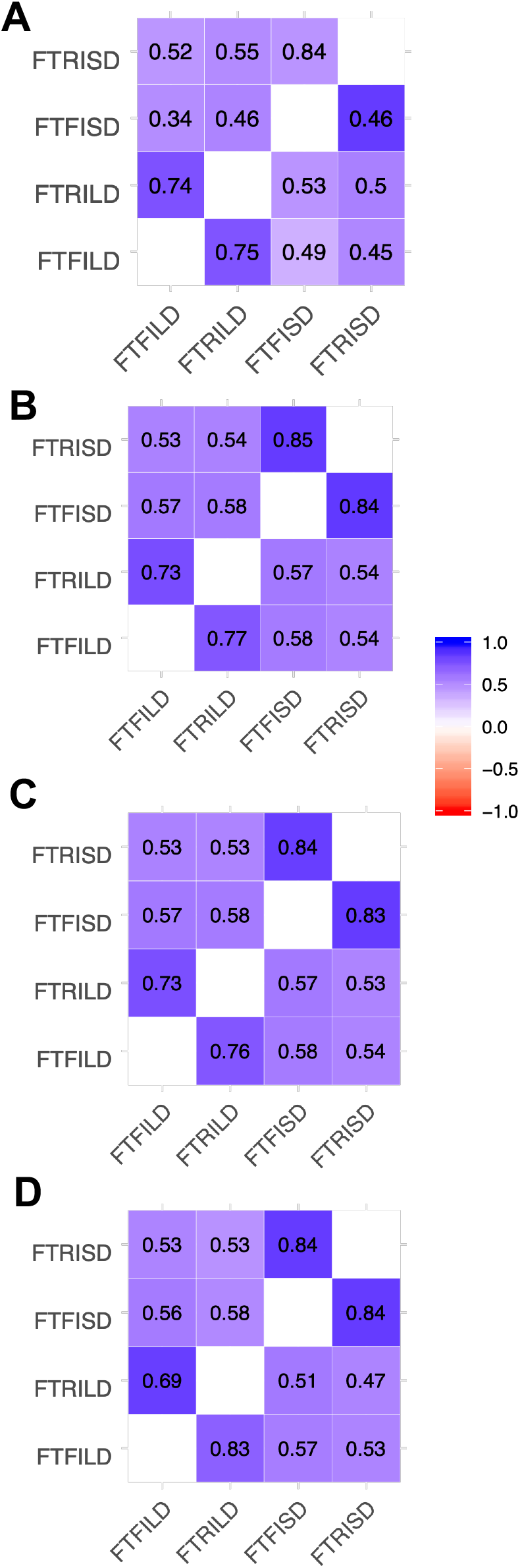
Environment by environment prediction values across GS models. Boxplots in each panel showed the distribution of intercept values across 100 cycles for (A) FxRRBLUP (Ridge Regression Best Linear Unbiased Prediction: Parametric model with fixed effects), (B) RKHS (Reproducing Kernel Hilbert Space; Semi-Parametric model), (C) RRBLUP (Ridge Regression Best Linear Unbiased Prediction: Parametric model with no fixed effects), and (D) SVR (Support Vector Regression: Non-Parametric model) for flowering time under full irrigation and long day (FTFILD), flowering time under restricted irrigation and long day (FTRILD), flowering time under full irrigation and short day (FTFISD), flowering time under restricted irrigation and short day (FTRISD)

**Figure S 15:**
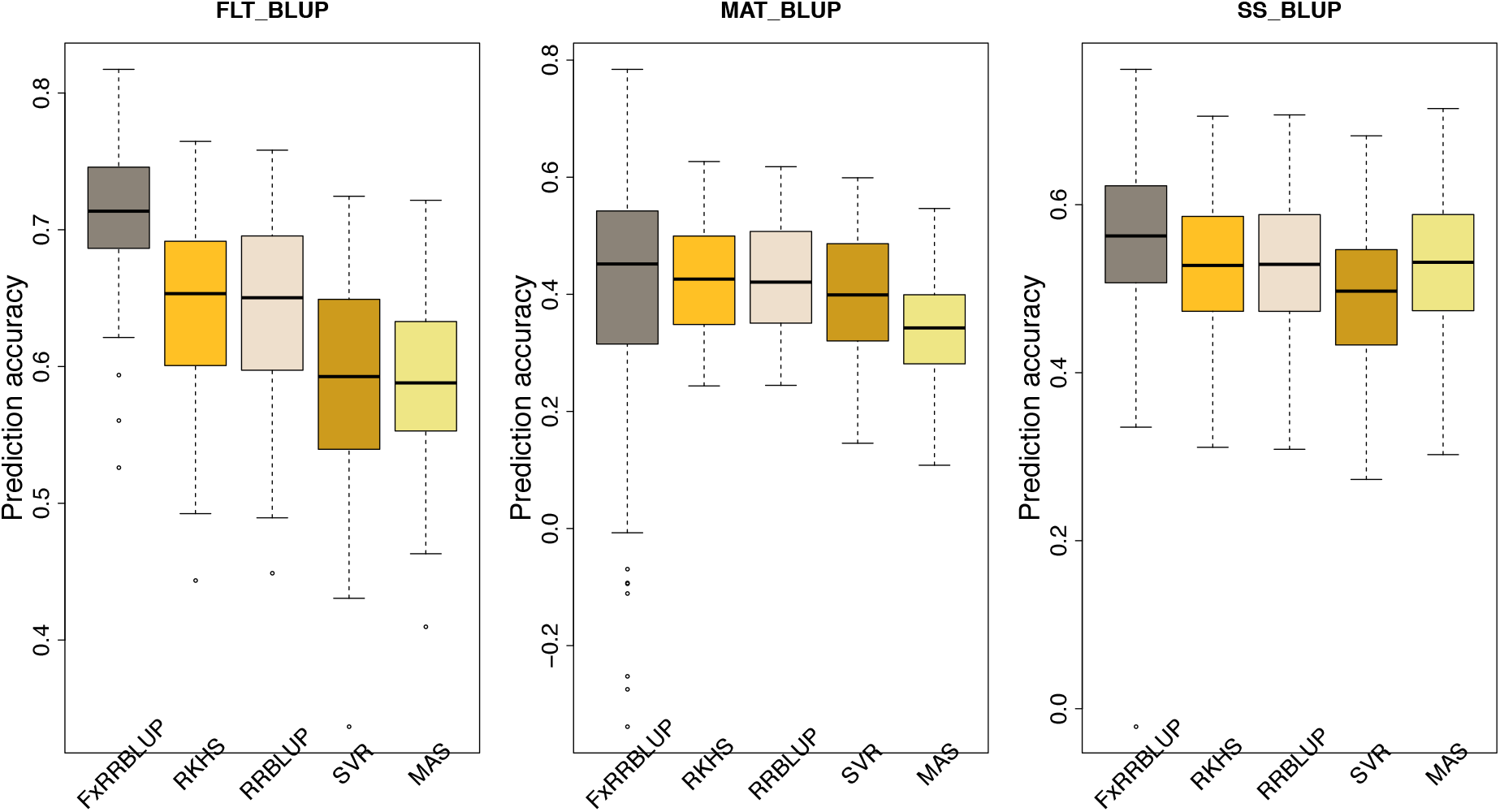
Comparison of prediction accuracy across GS and MAS models. Boxplots in each panel showed the distribution of prediction accuracy values across 100 cycles for FxRRBLUP (Ridge Regression Best Linear Unbiased Prediction: Parametric model with fixed effects), RKHS (Reproducing Kernel Hilbert Space; Semi-Parametric model), RRBLUP (Ridge Regression Best Linear Unbiased Prediction: Parametric model with no fixed effects), SVR (Support Vector Regression: Non-Parametric model), and MAS (Marker Assisted Selection) for flowering time BLUP (FLT_BLUP), maturity BLUP (MAT_BLUP), seed size BLUP (SS_BLUP).

**Figure S 16:**
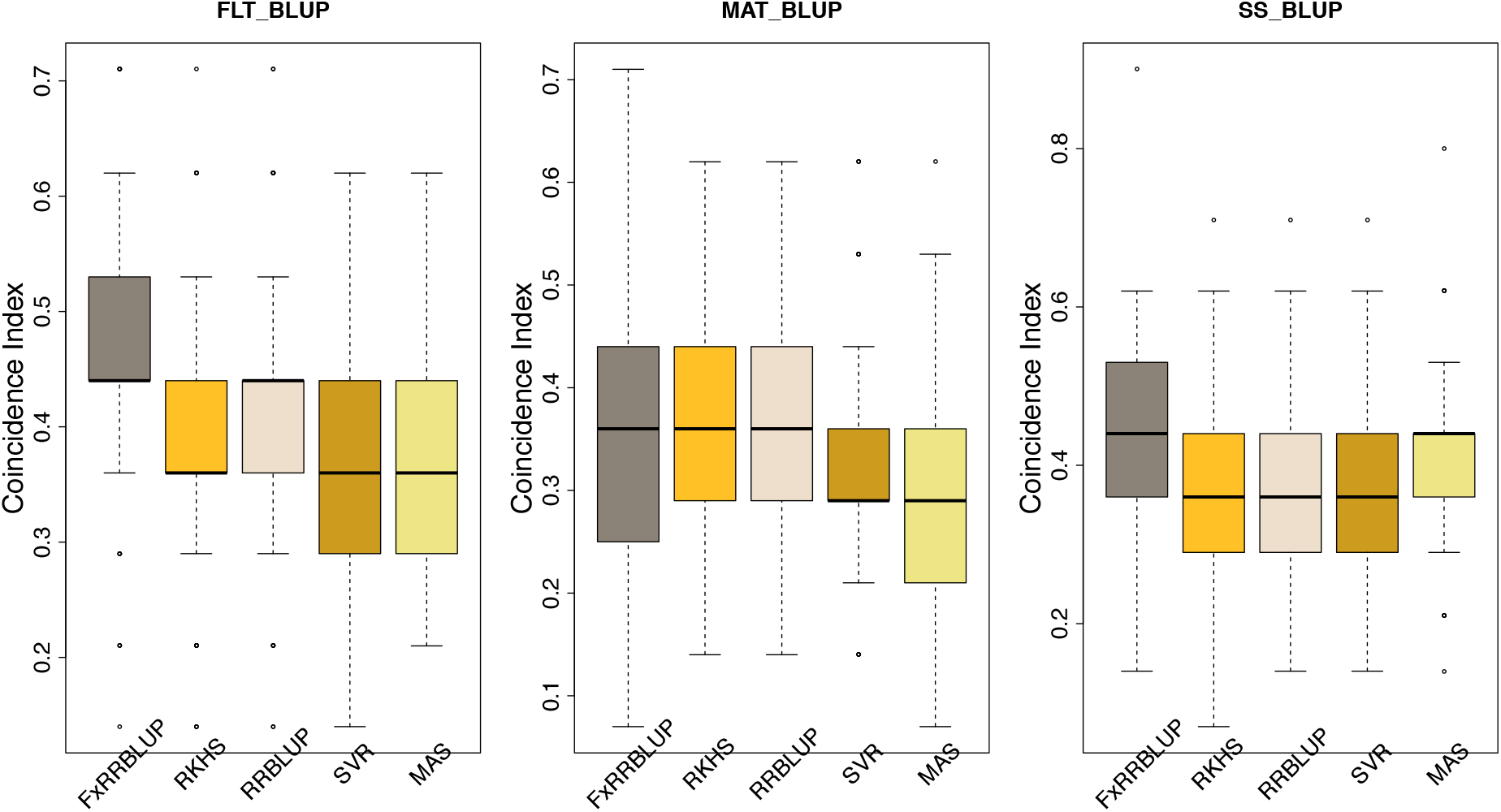
Comparison of coincidence index across GS and MAS models. Boxplots in each panel showed the distribution of coincidence index values across 100 cycles for FxRRBLUP (Ridge Regression Best Linear Unbiased Prediction: Parametric model with fixed effects), RKHS (Reproducing Kernel Hilbert Space; Semi-Parametric model), RRBLUP (Ridge Regression Best Linear Unbiased Prediction: Parametric model with no fixed effects), SVR (Support Vector Regression: Non-Parametric model), and MAS (Marker Assisted Selection) for flowering time BLUP (FLT_BLUP), maturity BLUP (MAT_BLUP), seed size BLUP (SS_BLUP).

**Figure S 17:**
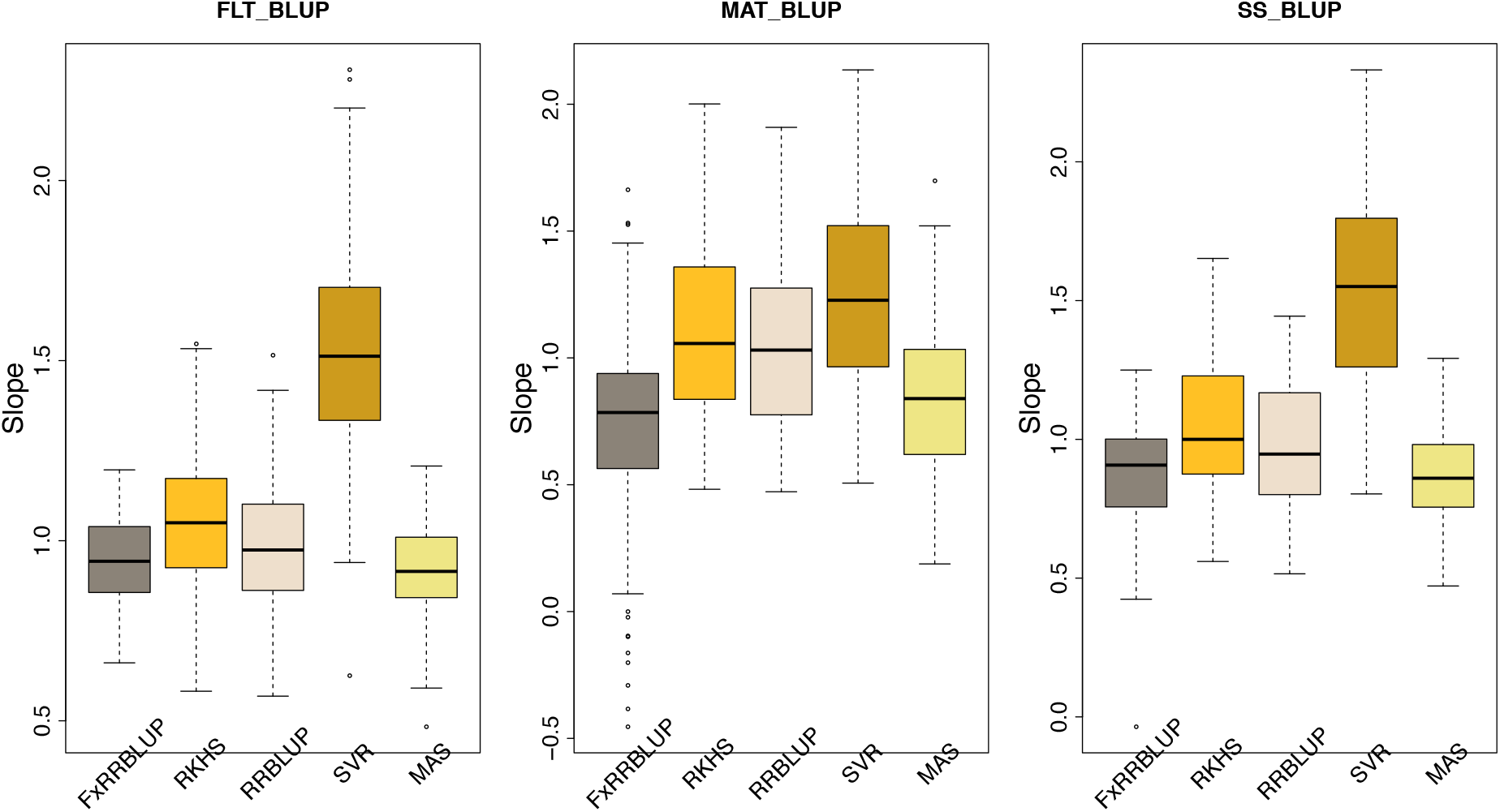
Comparison of slope values across GS and MAS models. Boxplots in each panel showed the distribution of slope values across 100 cycles for FxRRBLUP (Ridge Regression Best Linear Unbiased Prediction: Parametric model with fixed effects), RKHS (Reproducing Kernel Hilbert Space; Semi-Parametric model), RRBLUP (Ridge Regression Best Linear Unbiased Prediction: Parametric model with no fixed effects), SVR (Support Vector Regression: Non-Parametric model), and MAS (Marker Assisted Selection) for flowering time BLUP (FLT_BLUP), maturity BLUP (MAT_BLUP), seed size BLUP (SS_BLUP).

**Figure S 18:**
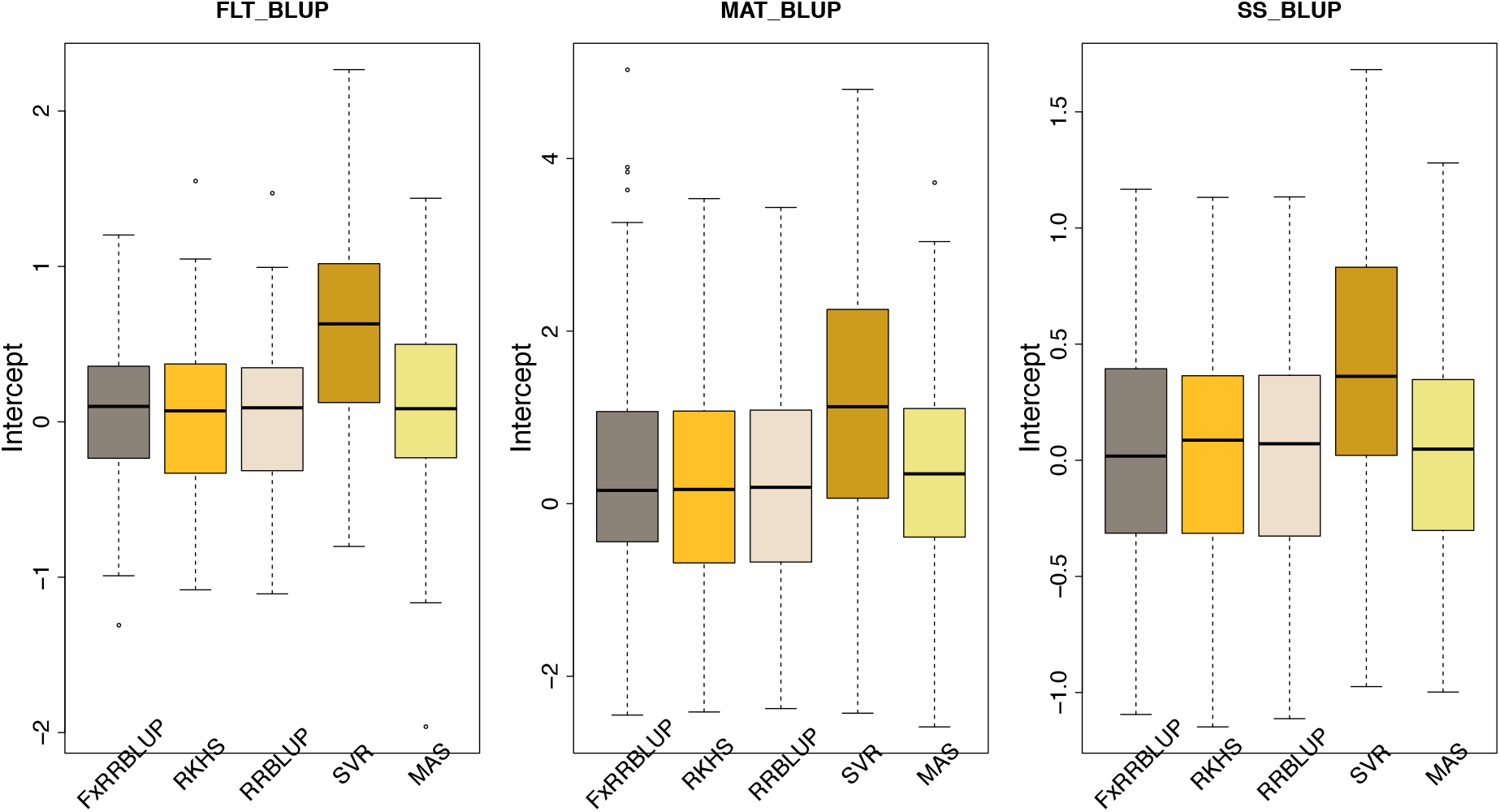
Comparison of intercept values across GS and MAS models. Boxplots in each panel showed the distribution of intercept values across 100 cycles for FxRRBLUP (Ridge Regression Best Linear Unbiased Prediction: Parametric model with fixed effects), RKHS (Reproducing Kernel Hilbert Space; Semi-Parametric model), RRBLUP (Ridge Regression Best Linear Unbiased Prediction: Parametric model with no fixed effects), SVR (Support Vector Regression: Non-Parametric model), and MAS (Marker Assisted Selection) for flowering time BLUP (FLT_BLUP), maturity BLUP (MAT_BLUP), seed size BLUP (SS_BLUP).

**Figure S 19:**
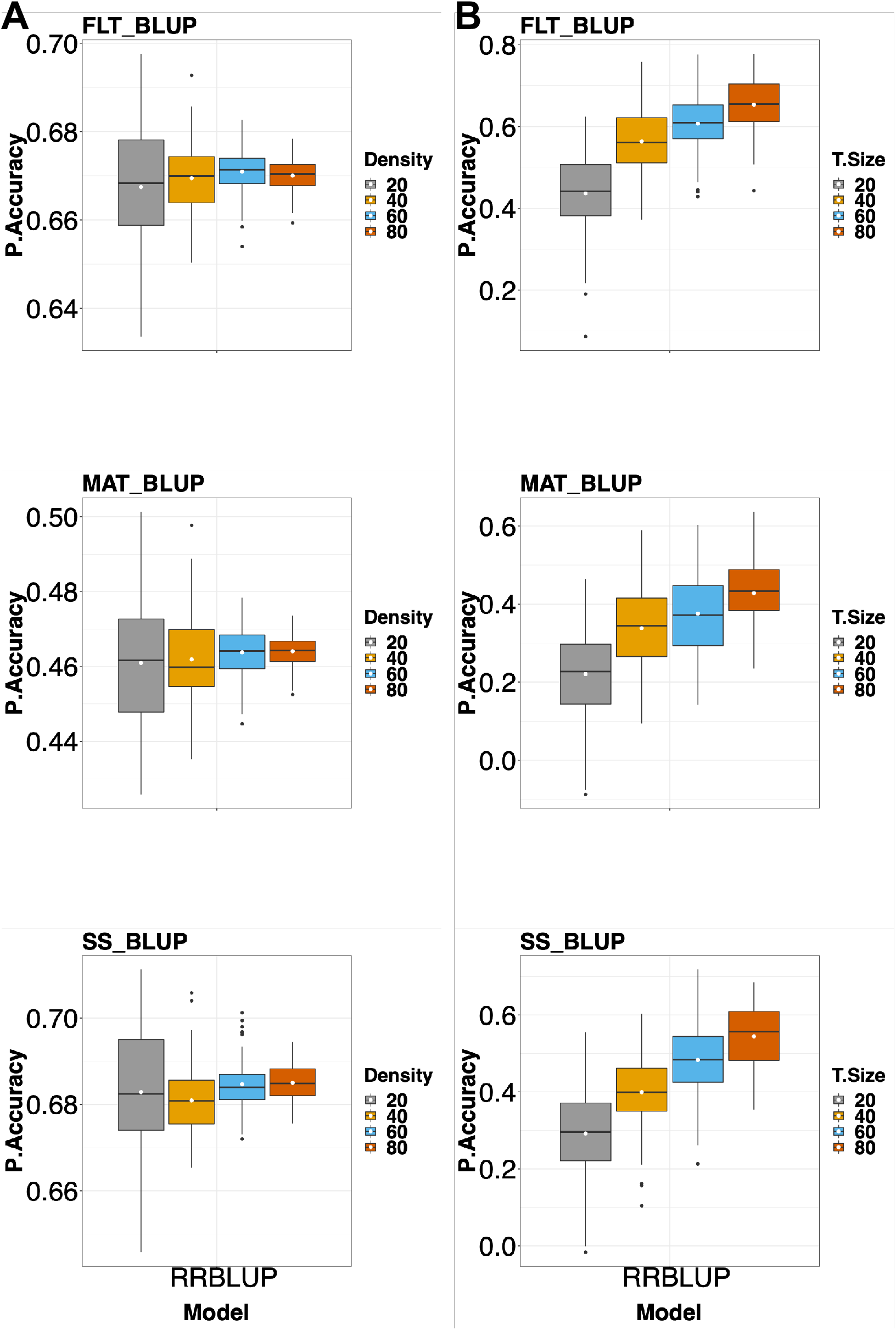
The effect of marker density and training population size on prediction accuracy. (A) Boxplots showing comparison among different marker densities (20%, 40%, 60%, and 80%). (B) Boxplots showing comparison among different training population sizes (20%, 40%, 60%, and 80%).

### Supplementary Tables

**Table S 1:**
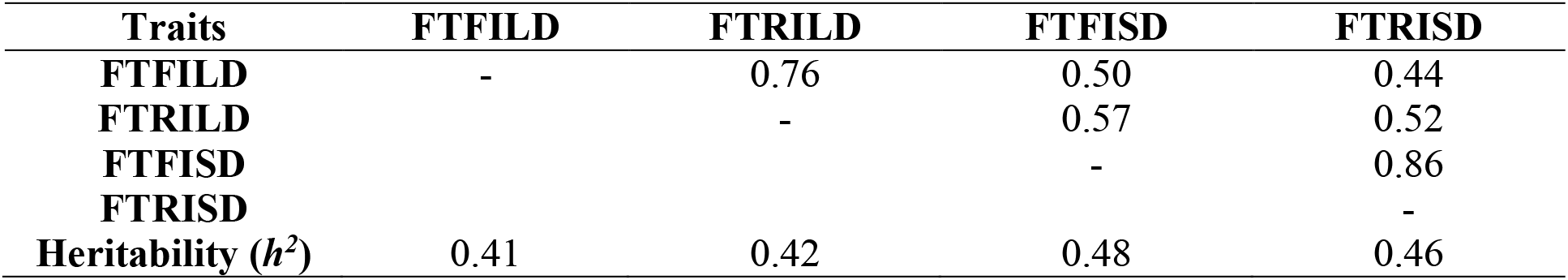
Environment by Environment correlation for flowering time.

**Table S 2:** Environment by environment correlation for maturity and seed size.

| Traits | MFISD | MRISD | SSFISD | SSRISD |
| --- | --- | --- | --- | --- |
| **Heritability ($h^2$)** | 0.30 | 0.21 | 0.47 | 0.39 |
| **MFISD** | - | 0.64 | - | - |
| **SSFISD** |  |  | - | 0.80 |

**Table S 3:**
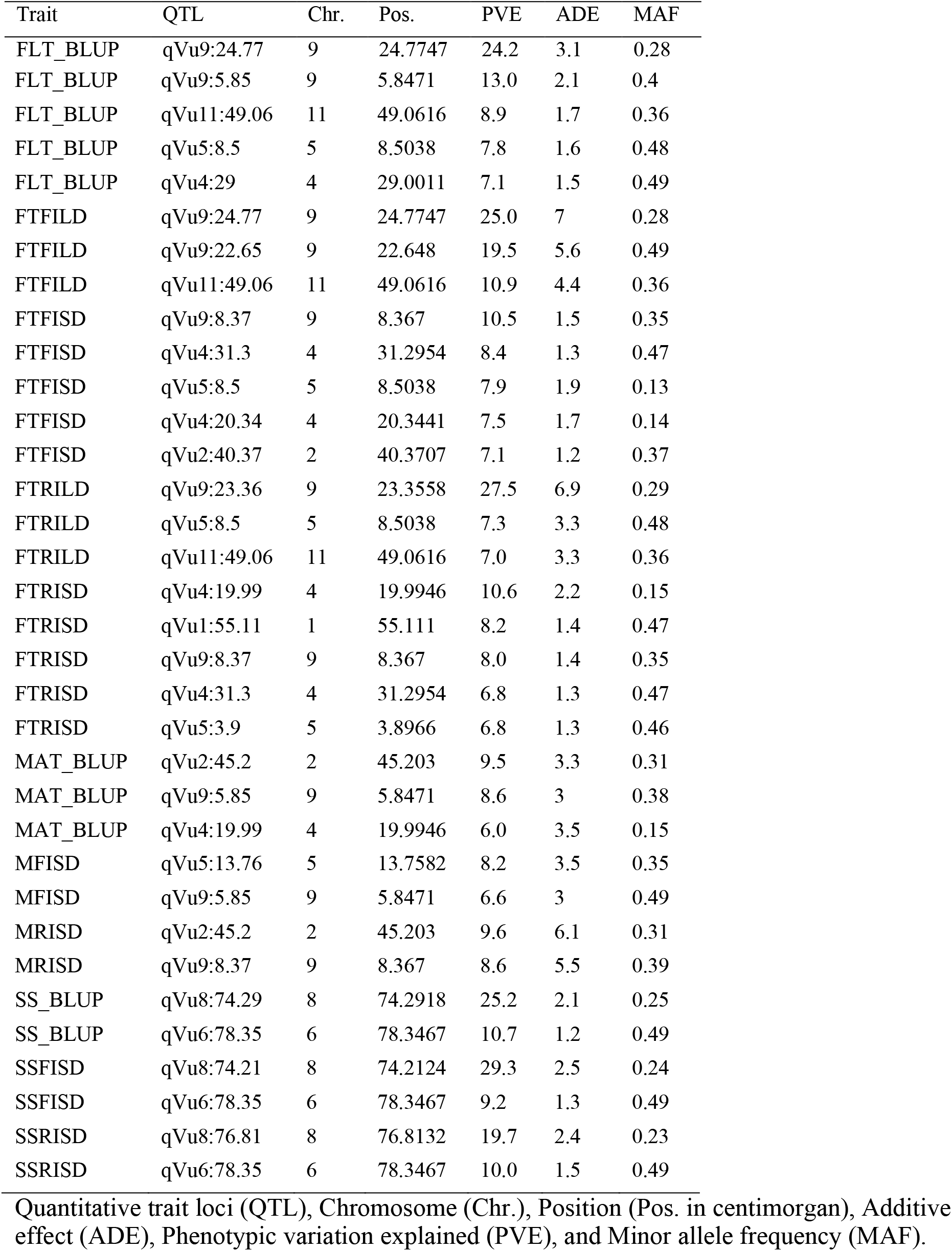
QTL identified by stepwise regression explaining at least 6% of phenotypic variation.

**Table S 4:** Epistatic QTL identified by SPAEML and their effect sizes.

| **Trait** | **QTL1** | **ADE1** | **MAF1** | **QTL2** | **ADE2** | **MAF2** |
| --- | --- | --- | --- | --- | --- | --- |
| FLT_BLUP | qVu9:25.39 | 3.1 | 0.28 | qVu11:62.84 | 1.9 | 0.14 |
| FLT_BLUP | qVu9:5.86 | 2.1 | 0.39 | qVu11:42.83 | 1.8 | 0.14 |
| FLT_BLUP | qVu5:12.79 | 1.7 | 0.34 | qVu6:78.36 | 0.5 | 0.47 |
| FLT_BLUP | qVu4:31.3 | 1.3 | 0.48 | qVu6:1.47 | 1.6 | 0.15 |
| FLT_BLUP | qVu1:66.57 | 1.6 | 0.12 | qVu9:26.8 | 1.5 | 0.26 |
| FLT_BLUP | qVu4:30.21 | 0.2 | 0.44 | qVu7:45.81 | 0.4 | 0.43 |
| FTFILD | qVu9:25.39 | 7 | 0.28 | qVu11:50.94 | 4.6 | 0.26 |
| FTFILD | qVu5:12.79 | 2.9 | 0.34 | qVu11:35.28 | 1.6 | 0.40 |
| FTFILD | qVu1:66.38 | 3.3 | 0.22 | qVu4:31.03 | 2.3 | 0.47 |
| FTFILD | qVu1:66.57 | 4.3 | 0.12 | qVu5:52.97 | 0.9 | 0.37 |
| FTFILD | qVu6:32.5 | 1.2 | 0.12 | qVu9:86.49 | 1.1 | 0.34 |
| FTFISD | qVu4:31.3 | 1.2 | 0.48 | qVu9:28.65 | 0.8 | 0.41 |
| FTFISD | qVu1:55.11 | 1.2 | 0.37 | qVu5:25.01 | 0.4 | 0.10 |
| FTFISD | qVu2:48.05 | 0.9 | 0.30 | qVu9:8.37 | 1.4 | 0.39 |
| FTFISD | qVu7:84.88 | 0.6 | 0.38 | qVu10:10.07 | 0.8 | 0.36 |
| FTFISD | qVu5:5.81 | 0.6 | 0.16 | qVu5:8.91 | 1 | 0.49 |
| FTRILD | qVu9:25.39 | 6.6 | 0.28 | qVu11:62.84 | 3.8 | 0.14 |
| FTRILD | qVu5:12.79 | 3.7 | 0.34 | qVu6:0.68 | 2.4 | 0.39 |
| FTRISD | qVu4:20.34 | 2.1 | 0.14 | qVu9:8.37 | 1.3 | 0.39 |
| FTRISD | qVu1:54.81 | 1.3 | 0.37 | qVu4:31.3 | 1.2 | 0.48 |
| FTRISD | qVu1:66.38 | 1.2 | 0.22 | qVu11:49.06 | 1.1 | 0.39 |
| FTRISD | qVu7:80.11 | 0.3 | 0.47 | qVu8:37.41 | 0.6 | 0.48 |
| MAT_BLUP | qVu2:48.05 | 2.8 | 0.30 | qVu9:8.37 | 3 | 0.39 |
| MFISD | qVu5:12.79 | 3.4 | 0.34 | qVu7:50.14 | 2 | 0.27 |
| MFISD | qVu2:39.29 | 2.4 | 0.36 | qVu8:73.15 | 2.4 | 0.34 |
| MFISD | qVu5:8.5 | 2.8 | 0.19 | qVu9:8.37 | 2.8 | 0.39 |
| MRISD | qVu2:48.05 | 5.3 | 0.30 | qVu9:8.37 | 5.5 | 0.39 |
| MRISD | qVu2:35.19 | 5.4 | 0.21 | qVu8:75.88 | 4.8 | 0.24 |
| SS_BLUP | qVu6:78.36 | 1.1 | 0.47 | qVu7:18.1 | 0.4 | 0.11 |
| SS_BLUP | qVu6:78.35 | 0.8 | 0.13 | qVu8:74.29 | 2.1 | 0.25 |
| SSFISD | qVu4:62.75 | 0.8 | 0.10 | qVu8:74.29 | 2.5 | 0.25 |
| SSFISD | qVu5:20.54 | 0.6 | 0.11 | qVu6:78.36 | 1.2 | 0.47 |
| SSRISD | qVu6:78.35 | 1 | 0.13 | qVu8:74.29 | 2.2 | 0.25 |
| SSRISD | qVu6:3.67 | 0.8 | 0.12 | qVu6:78.36 | 1.4 | 0.47 |

Quantitative trait loci (QTL), Linkage group (LG), Position (Pos. in centimorgan), Additive effect (ADE), , and Minor allele frequency (MAF).

**Table S 5:** Mean and standard deviation of prediction accuracy across GS and MAS models.

| Trait | FxRRBLUP | RKHS | RRBLUP | SVR | MAS |
| --- | --- | --- | --- | --- | --- |
| FT_BLUP | 0.71±0.05 | 0.65±0.07 | 0.65±0.07 | 0.59±0.08 | 0.59±0.06 |
| FTFILD | 0.68±0.05 | 0.55±0.08 | 0.55±0.08 | 0.54±0.08 | 0.64±0.06 |
| FTFISD | 0.56±0.20 | 0.59±0.07 | 0.59±0.07 | 0.55±0.08 | 0.33±0.11 |
| FTRILD | 0.68±0.07 | 0.58±0.08 | 0.58±0.07 | 0.50±0.08 | 0.61±0.08 |
| FTRISD | 0.58±0.17 | 0.58±0.07 | 0.58±0.07 | 0.53±0.08 | 0.25±0.10 |
| MT_BLUP | 0.40±0.23 | 0.42±0.09 | 0.42±0.09 | 0.40±0.10 | 0.33±0.09 |
| MFISD | 0.30±0.20 | 0.39±0.09 | 0.39±0.09 | 0.36±0.09 | 0.20±0.10 |
| MRISD | 0.25±0.29 | 0.37±0.11 | 0.37±0.11 | 0.34±0.12 | 0.30±0.10 |
| SS_BLUP | 0.56±0.10 | 0.52±0.08 | 0.53±0.08 | 0.49±0.09 | 0.53±0.09 |
| SSFISD | 0.58±0.14 | 0.54±0.08 | 0.54±0.08 | 0.51±0.09 | 0.57±0.08 |
| SSRISD | 0.50±0.11 | 0.45±0.08 | 0.45±0.08 | 0.43±0.09 | 0.47±0.10 |

**Table S 6:** Mean and standard deviation of coincidence index of GS and MAS models.

| Trait | FxRRBLUP | RKHS | RRBLUP | SVR | MAS |
| --- | --- | --- | --- | --- | --- |
| FLT_BLUP | 0.47±0.11 | 0.40±0.11 | 0.42±0.11 | 0.37±0.10 | 0.37±0.09 |
| FTFILD | 0.49±0.09 | 0.37±0.10 | 0.37±0.10 | 0.35±0.11 | 0.45±0.10 |
| FTFISD | 0.43±0.16 | 0.44±0.10 | 0.44±0.10 | 0.42±0.09 | 0.30±0.11 |
| FTRILD | 0.40±0.09 | 0.36±0.10 | 0.35±0.10 | 0.28±0.10 | 0.37±0.10 |
| FTRISD | 0.45±0.15 | 0.43±0.09 | 0.42±0.10 | 0.42±0.10 | 0.26±0.09 |
| MT_BLUP | 0.34±0.15 | 0.35±0.09 | 0.36±0.10 | 0.33±0.11 | 0.27±0.10 |
| MFISD | 0.32±0.12 | 0.30±0.09 | 0.30±0.10 | 0.31±0.09 | 0.26±0.09 |
| MRISD | 0.31±0.18 | 0.34±0.10 | 0.35±0.10 | 0.33±0.10 | 0.33±0.10 |
| SS_BLUP | 0.44±0.11 | 0.37±0.11 | 0.36±0.10 | 0.36±0.10 | 0.42±0.11 |
| SSFISD | 0.48±0.11 | 0.44±0.09 | 0.44±0.09 | 0.43±0.09 | 0.46±0.10 |
| SSRISD | 0.42±0.11 | 0.36±0.11 | 0.37±0.11 | 0.36±0.10 | 0.40±0.10 |

